# The spatial Muller’s ratchet: surfing of deleterious mutations during range expansion

**DOI:** 10.1101/2019.12.20.883975

**Authors:** Félix Foutel-Rodier, Alison Etheridge

## Abstract

During a range expansion, deleterious mutations can “surf” on the colonisation front. The resultant decrease in fitness is known as expansion load. An Allee effect is known to reduce the loss of genetic diversity of expanding populations, by changing the nature of the expansion from “pulled” to “pushed”. We study the impact of an Allee effect on the formation of an expansion load with a new model, in which individuals have the genetic structure of a Muller’s ratchet. A key feature of Muller’s ratchet is that the population fatally accumulates deleterious mutations due to the stochastic loss of the fittest individuals, an event called a click of the ratchet. We observe fast clicks of the ratchet at the colonization front owing to small population size, followed by a slow fitness recovery due to migration of fit individuals from the bulk of the population, leading to a transient expansion load. For large population size, we are able to derive quantitative features of the expansion wave, such as the wave speed and the frequency of individuals carrying a given number of mutations. Using simulations, we show that the presence of an Allee effect reduces the rate at which clicks occur at the front, and thus reduces the expansion load.

## 1 Introduction

### Gene surfing and expansion load

The genetics of range expansion is a complex topic that has attracted much attention. In a pioneering work, [10] reported that during a range expansion a neutral mutant appearing in the front of an expansion could rapidly spread over a vast region of space. This phenomenon was further studied in [31] and dubbed gene surfing, see Figure 1 for an illustration. Gene surfing originates from two features of range expansions. First, the population density is lower at the range’s margin than in its core, where the population has had more time to grow to carrying capacity. Thus a mutant that appears there is already in relatively high frequency among the few individuals in the front. Moreover, individual-level demographic stochasticity, which is the cause of population-level genetic drift, can lead to a further rapid increase of the local frequency of this mutant. Second, population spread can be caricatured by successive founding events, where a few individuals migrate to an empty habitat and grow a new subpopulation. Individuals living at the edge are more likely to be recruited to found these new subpopulations as they are spatially closer to the empty habitats. In other words, individuals that form the subsequent front are sampled from the current front, not from the bulk. Combining these two features, the initial increase in frequency of the mutant at the front (which is the result of the small population size) gets amplified by the successive resampling from the front, and the mutant can reach a high frequency over a large spatial area, see Figure 1, panel (c). Gene surfing is now a well-understood phenomenon, that has been assessed both theoretically [10, 31, 48, 26], and empirically using microbial growth experiments [28, 27] or naturally occuring genetic data [22]. See [12, 13] for reviews of this topic.

**Figure 1:**
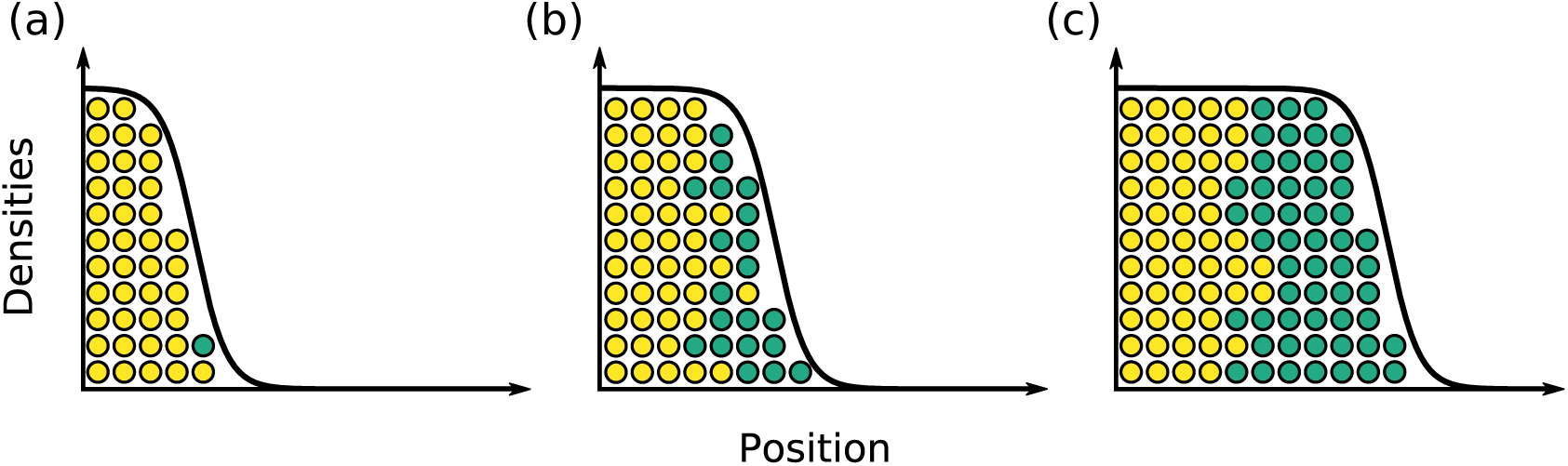
Illustration of gene surfing. (a) a mutant (in green) appears in the front of an expanding population; (b) the mutant rapidly reaches fixation in the front; (c) the mutant offspring further expand, leading to a clear allele segregation.

The very first step of the surfing phenomenon is the local increase in frequency of an allele at the front, resulting from the increased growth rate when population density is low. Increased genetic drift at the front makes reaching fixation easier for both neutral and deleterious mutants. Hence it is not surprising that deleterious mutations can also surf [48]. As deleterious mutations are more frequent than beneficial ones [14], it has been predicted and assessed in [39] that fitness at the front is decreasing during a range expansion, due to successive surfing of deleterious mutations along the expansion axis. This reduction in fitness due to range expansion is known as *expansion load*. Expansion load is thus the additional fitness disadvantage that a population has accumulated during a range expansion, due to a reduced ability of selection to purge deleterious mutations, see [41] for a review. While expansion load has been clearly highlighted using simulations [39, 38, 40, 19], genomic evidence of an expansion load remains a debated topic [8, 44]. The presence of an expansion load has been reported in human populations after the Out of Africa expansion [29, 42], and in plants [21, 50], see [41] for a review.

### Impact of an Allee effect

A population exhibits an Allee effect if its maximal per-capita growth rate is achieved at intermediate population density rather than at low population density. The Allee effect is said to be *strong* if the per-capita growth rate is negative at low population density, that is if the population is unable to grow under a certain critical density threshold [7, 47]. Otherwise the Allee effect is *weak*. Allee effects arise in many biological contexts, including for example the presence of cooperation, or the difficulty in finding mates for reproduction [34, 45].

In the context of range expansion, an Allee effect shifts the location of the individuals with the highest per-capita growth rate towards the bulk of the population, see Figure 2 panel (a). In the absence of an Allee effect, individuals at the leading edge of the front, where the population density is the lowest, have the highest growth rate. The wave is *pulled* by these individuals in the front. Conversely, if the Allee effect is strong enough, individuals with the highest growth rate are located midway between the front and the bulk of the population. The wave is then *pushed* by the bulk of the population. The distinction between pushed and pulled waves is a well-established paradigm of the reaction-diffusion literature, see [46, 49]. The correspondence between the pulled/pushed nature of the waves and the absence/presence of an Allee effect is not perfect. A weak Allee effect can lead to both pulled and pushed waves, as is for instance the case in the model considered in [5]. Nevertheless, an Allee effect is a necessary condition for the wave to be pushed [46], and a strong Allee effect is a sufficient one.

**Figure 2:**
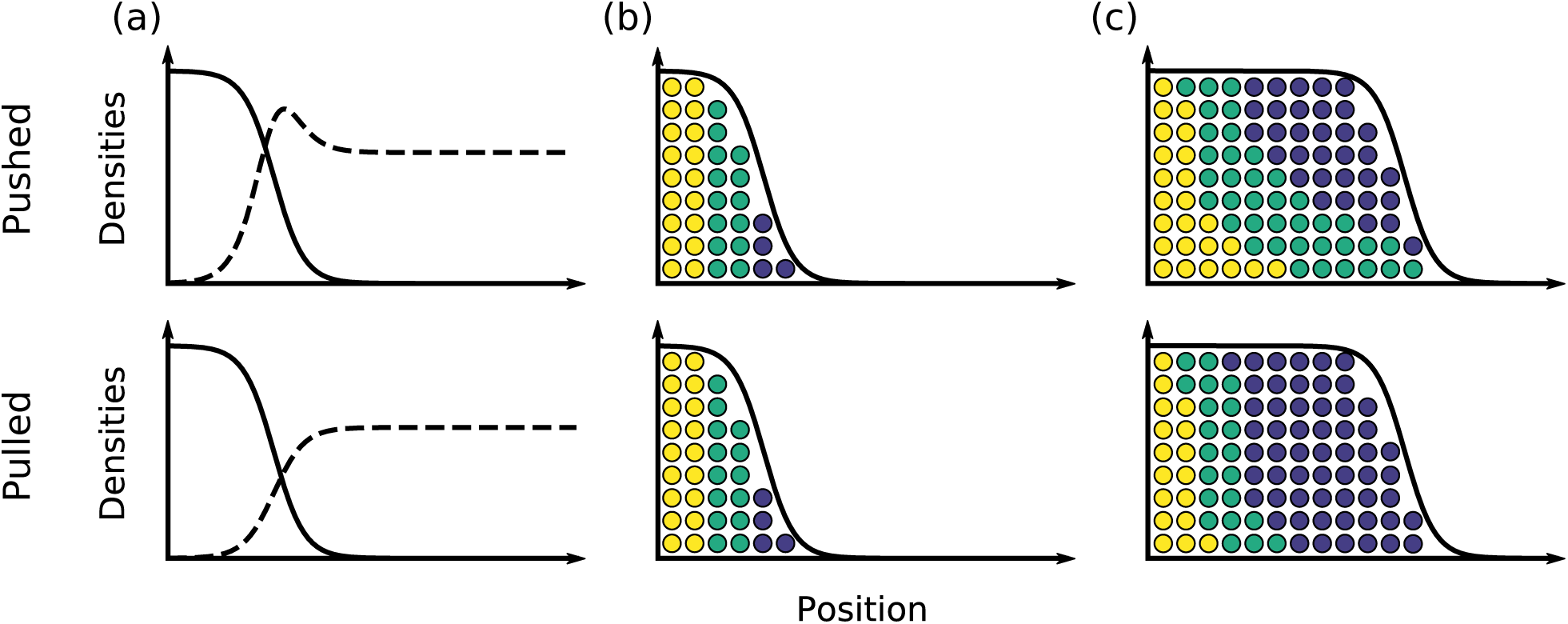
Illustration of the impact of an Allee effect. (a) Solid line. population density; dashed line. per-capita growth rate. (b) Particles have been labeled in different colours according to their initial location. (c) Genetic composition of the population after expansion. In the pulled wave, only the few individuals in the front are the founders of the new habitats, while in the pushed wave individuals from the bulk also contribute to the new front.

From a genetic perspective, an Allee effect increases the “effective population size” of the front. Individuals that leave the largest number of offspring are either located in the front in the pulled case, or towards the bulk in the pushed case. Thus, we expect that only the very few individuals far in the front contribute to the genetic pool of the population in a pulled wave, leading to a more drastic loss of diversity than in a pushed wave, see Figure 2 for an illustration. Several theoretical studies have assessed the impact of an Allee effect on the genetics of range expansion [43, 18, 26, 5] in a neutral setting, let us briefly review their results.

The authors of [26] considered a stochastic particle system modelling range expansion, and studied the fixation probability of individuals in the front as a function of their locations. Using both simulations and analytical approximations, they provided an expression for the fixation probability, and showed that it reaches a maximum at a location which is shifting towards the bulk of the population as the strength of the Allee effect increases. In [43, 18] the authors considered a deterministic partial differential equation analogous to the celebrated Fisher-KPP equation [17, 32], which has been widely used to model invading populations. They divided the total population into several neutral fractions, and studied the long-term fate of these fractions. They proved that in the pulled case, only the fraction closest to the front is able to follow the expansion, and that all other fractions are left behind. In the long run, the population is only composed of the offspring of the individuals that were initially closest to the front. Conversely in the pushed case all fractions are able to follow the expansion. Asymptotically, all individuals in the population leave progeny that live in the front. The long-term contribution to the front of the various initial fractions can be computed explicitly, with an expression consistent with the approximate fixation probability found in [26]. Finally, [5] used simulations and analytical approximations to study the rate of loss of genetic diversity during a range expansion. They showed that, as the strength of the Allee effect increases, the loss of genetic diversity is slowed down.

All the above studies consistently find that an Allee effect impedes gene surfing, and rescues the genetic diversity in the front of an expanding population. Expansion load originates from successive surfing of deleterious mutants, we thus expect the presence of an Allee effect to reduce expansion load. Nevertheless, the impact of an Allee effect on expansion load has never been explicitly tested. All existing theoretical studies on expansion load assume logistic growth of the population, in particular no Allee effect [39, 40, 38]. Moreover, the only analytical results available for these models have been derived using a serial founder approximation. In this approximation, the front is supposed to be genetically isolated from the bulk, and at each time-step, a new front is formed by sampling a few individuals from the previous front and letting them grow logistically. Even if this approach yields good approximations of the mean fitness at the front, it misses the continuous gene flow between the bulk and the front that occurs during range expansion, especially in the presence of an Allee effect. The objectives of the present work are thus twofold. First we aim to study the impact of an Allee effect on the expansion load. We will restrict our attention to the case of a weak Allee effect, and will not consider the impact of a strong Allee effect. Second, we aim to build a model that is more amenable to continuous space techniques, in order to take into account the entire dynamics of the expanding population.

### A spatial Muller’s ratchet

In order to keep the genetic structure of the population as simple as possible, we consider genetic dynamics similar to that of [25], leading to a Muller’s ratchet [16, 37]. Muller’s ratchet is a mechanism that was first proposed as an explanation for the evolution of recombination, it can be formulated as follows. Consider a population of finite size that can only accumulate deleterious mutations over time. If mutations are irreversible and negatively selected, without drift the population should reach a mutation-selection equilibrium. Nevertheless, due to genetic drift all individuals without mutations are eventually lost. At such a time, in the absence of recombination, the minimal number of mutations in the population is permanently increased by one. We say that the ratchet has clicked. At each click of the ratchet the fitness of the population is decreased and successive clicks of the ratchet should drive it towards extinction. In the presence of recombination, chromosomes without mutations can be recreated by recombining two chromosomes with mutations at different loci, rescuing the population from the ratchet, and thus giving recombination a selective advantage.

In our context, we consider an expanding population where individuals can only accumulate deleterious mutations, without possible reversion or recombination. Due to higher genetic drift at the front, we expect the ratchet to click more often in the front than in the bulk. After a click at the front, we expect the population to be separated into two distinct regions: one towards the front where the ratchet has clicked, and the other towards the bulk where the ratchet has not clicked. Despite their lower fitness, individuals in the front will still be able to colonise new habitat as the low population density guarantees a positive growth rate. Interestingly, individuals without mutations have a positive growth rate when they are placed in a location where the ratchet has clicked, as their fitness is larger. Therefore, the region where the ratchet has not clicked should also be able to expand into the region where it has, at a rate depending on the fitness difference between the two regions. This can be thought of as an inner expansion wave evolving inside the larger expansion wave of the whole population. Each click of the ratchet should create a new inner wave of less unfit individuals. Successive rapid clicks of the ratchet at the front will create an expansion load, but there will be some recovery of fitness at any given location as fitter individuals from the “bulk” invade. This phenomenon will be illustrated numerically in the forthcoming Section 3.2. Let us spell out the dynamics of the model more precisely.

## 2 Methods

### Description of the model

We consider a population of non-recombining individuals each carrying a single chromosome. The population is subdivided in demes indexed by ℤ. Each individual is entirely characterized by the number of deleterious mutations it carries and its spatial location. We record as *n*_*i,k*_(*t*) the number of individuals carrying *k* ≥ 0 mutations in deme *i* ∈ ℤ at time *t* ∈ ℝ_+_, and let *N*_*i*_(*t*) = ∑_*k*≥0_ *n*_*i,k*_(*t*) be the total population size in deme *i*. Thus the vector (*n*_*i,k*_(*t*); *i* ∈ ℤ, *k* ≥ 0) contains all the information about the population at time *t*.

Time is continuous and individuals can reproduce, die or migrate according to the following rules. An individual located in deme *i* and carrying *k* mutations gives birth to a new individual at rate *λ*_*k*_(*N*_*i*_), and dies at rate *δ*(*N*_*i*_), where

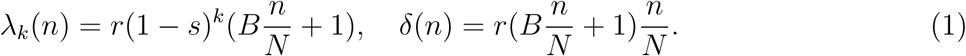

The offspring is located in the same deme as its parent. With probability 1 − *µ*, it inherits the same number of mutations as its parent, and with probability *µ*, it accumulates an additional one. Finally each individual migrates at rate *m*, and goes to one of the two nearest demes with equal probability.

Let us provide an intuitive description of equation (1) and of the parameters of the model. All deleterious mutations have the same fitness effect *s* > 0, and fitness is multiplicative across loci. If *B* = 0, we recover a stochastic version of the usual logistic growth, where *r* is the Malthusian growth parameter and *N* is a scaling parameter that can be thought of as the local carrying capacity of the population. Taking *B* > 0 introduces a density-dependence in the growth rate of the population. We think of *B* as a cooperation parameter, where cooperation acts on the overall growth rate of the population, which will tune the strength of the Allee effect. Notice that the function *n* ↦ *λ*_0_(*n*) − *δ*(*n*) is non-negative, and that it reaches its maximum for 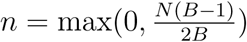. Thus, for *B* ≤ 1, the per-capita growth rate is maximal for *n* = 0, i.e., there is no Allee effect, while for *B* > 1, we see a weak Allee effect. Increasing *B* increases the strength of that Allee effect, as it further shifts the location of the maximal per-capita growth rate to higher population density. This parametrization of the Allee effect is similar to the one considered in [5].

### Large population scalings

As is usual in population genetics, in order to obtain analytical results about our model, we consider a large population size scaling. We begin with the deterministic infinite population limit. For a fixed value of *N*, let 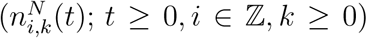 be a realization of the above model, with population size parameter *N* and migration rate *m*_*N*_. Let *L* be a space renormalization parameter, and for *i* ∈ ℤ and *x* = *i/L* set

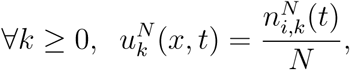

and interpolate the function 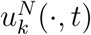 linearly between the points {*i/L* : *i* ∈ ℤ}. Then a standard generator calculation (see Appendix A.1) suggests that provided the initial condition converges, and *m*_*N*_ */L*^2^ → *m*, then, as 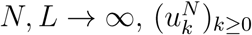 converges to the solution of

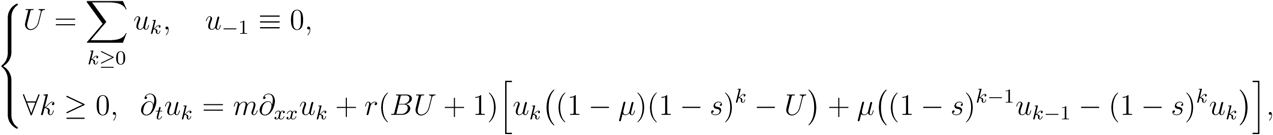

started from the corresponding initial condition. In what follows, we will always assume that selection is weak, and that mutation is low, i.e., that *s, µ* ≪ 1. To the first order in *s* and *µ*, the above equation becomes

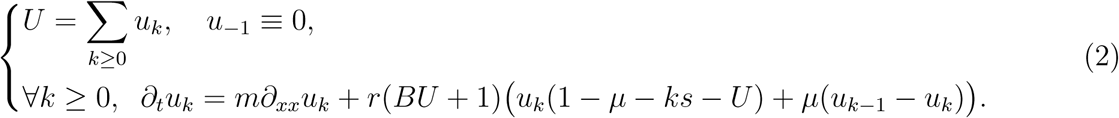

This limit is deterministic and thus does not take genetic drift into account and we do not observe gene surfing. By retaining terms up to order 1*/N* in a generator calculation, we can derive a diffusion approximation for our model that accounts for finite size fluctuations, see Appendix A.2. Under a diffusive scaling, where the population size is further rescaled by a factor *L*, and time is rescaled by a factor *L*^2^, when *N* is large, the process (*u*_*k*_)_*k*≥0_ is approximated by the following system of stochastic partial differential equations

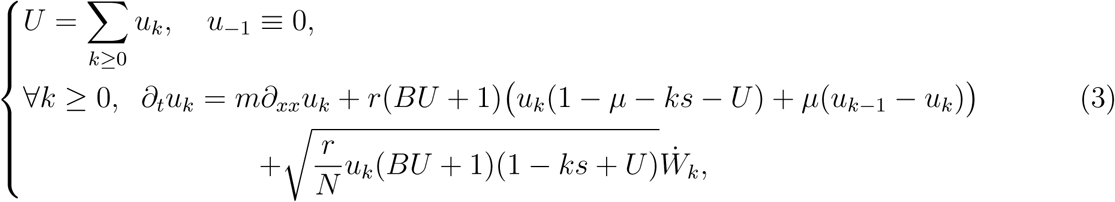

where 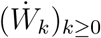 are independent space-time white noises.

The above two limits have been obtained by qualitative comparison of the generator of 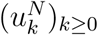 for large *N*. We do *not* prove the convergence of the process to any of these limiting objects, which is a highly non-trivial problem. Nevertheless, see [9, 36] for rigorous treatments of similar convergence results.

### Simulation setup

When started from a finite number of individuals, our model is a simple continuous-time Markov chain that we simulate using the following classical algorithm: at each iteration of the algorithm, we compute the total transition rate 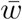 of the population and increment the time *t* by an exponential variable with mean 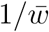. The transition that occurs is then chosen independently with probability proportional to the transition rates. Notice that *t* will always refer to the “actual time” of the simulation and *not* to the number of iterations. In each simulation, at *t* = 0 only the first 30 demes are occupied, all other demes are empty. The initial number of individuals in the occupied demes, and the distribution of the number of mutations, is chosen according to their deterministic equilibrium value, computed in (6). We restricted the spatial domain to 500 demes and used reflecting boundary conditions for the migration, i.e., individuals are not allowed to move outside the domain.

### Estimation of the click rate

In order to estimate the click rate, we need to determine from the simulation the moment when the ratchet has clicked. Let us denote by 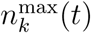 the location of the right-most deme containing individuals carrying *k* mutations at time *t*, defined as

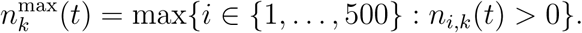

We define the approximate first click time *T*_1_ of the ratchet as

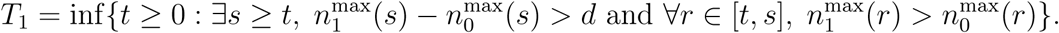

In words, *T*_1_ is the first moment when individuals with one mutation get ahead of individuals with no mutations, and will get *d* demes ahead before being caught up. We set *d* = 30 by observing from the simulations that once 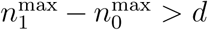, it is very unlikely that the inner wave catches up the front of expansion before the population has colonized the entire habitat.

In order to obtain the mean time to the first click, we averaged *T*_1_ over many simulations for various parameter values. Once the population has expanded, as there are many more individuals in the bulk than in the front, most of the events that occur are reproduction events in the bulk. These events are not relevant for the computation of the click time as individuals far in the bulk will never be able to reach the front. Thus in order to speed the simulations up, a frame of width *d*, co-moving with the front, was used for the simulations of Figure 5. We set the birth, death and migration rates of all individuals in deme *i* such that 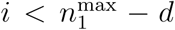 to 0, and also prevent individuals in deme 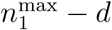 from migrating to the left. We emphasize that the co-moving frame was only used in the simulations of Figure 5 and that all other simulations account for the events that occur in the bulk.

**Figure 3:**
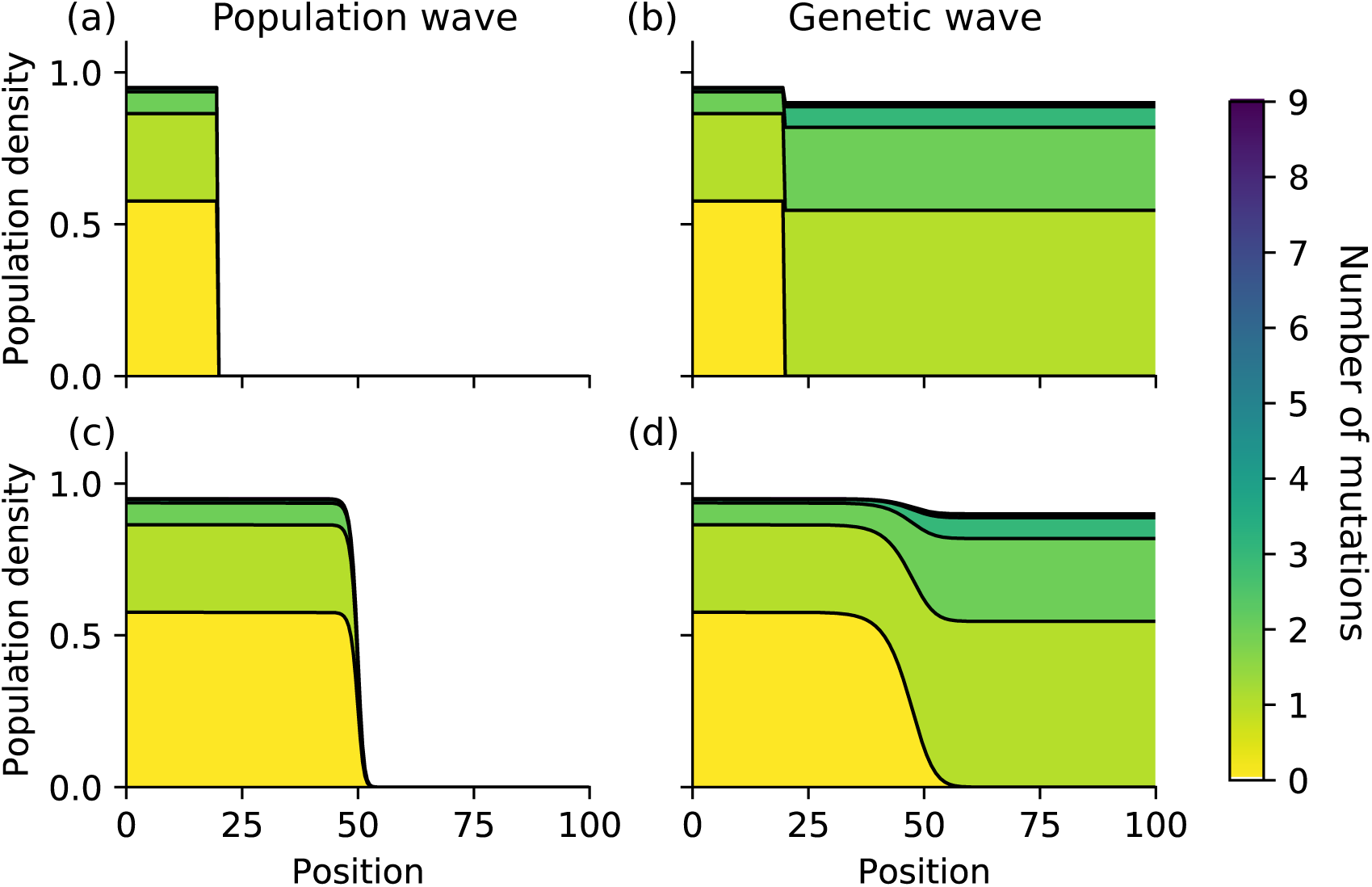
Simulation of (2) with different initial conditions. (a, c) Population travelling wave at *t* = 0 (a) and *t* = 50 (c). (b, d) Genetic travelling wave at *t* = 0 (b) and *t* = 250 (c); the initial condition is of the form (10) for *x*_0_ = 20. Parameter values are *r* = 1, m = 0.1; *s* = 0.05, *μ* = 0.025, *B* = 0.

**Figure 4:**
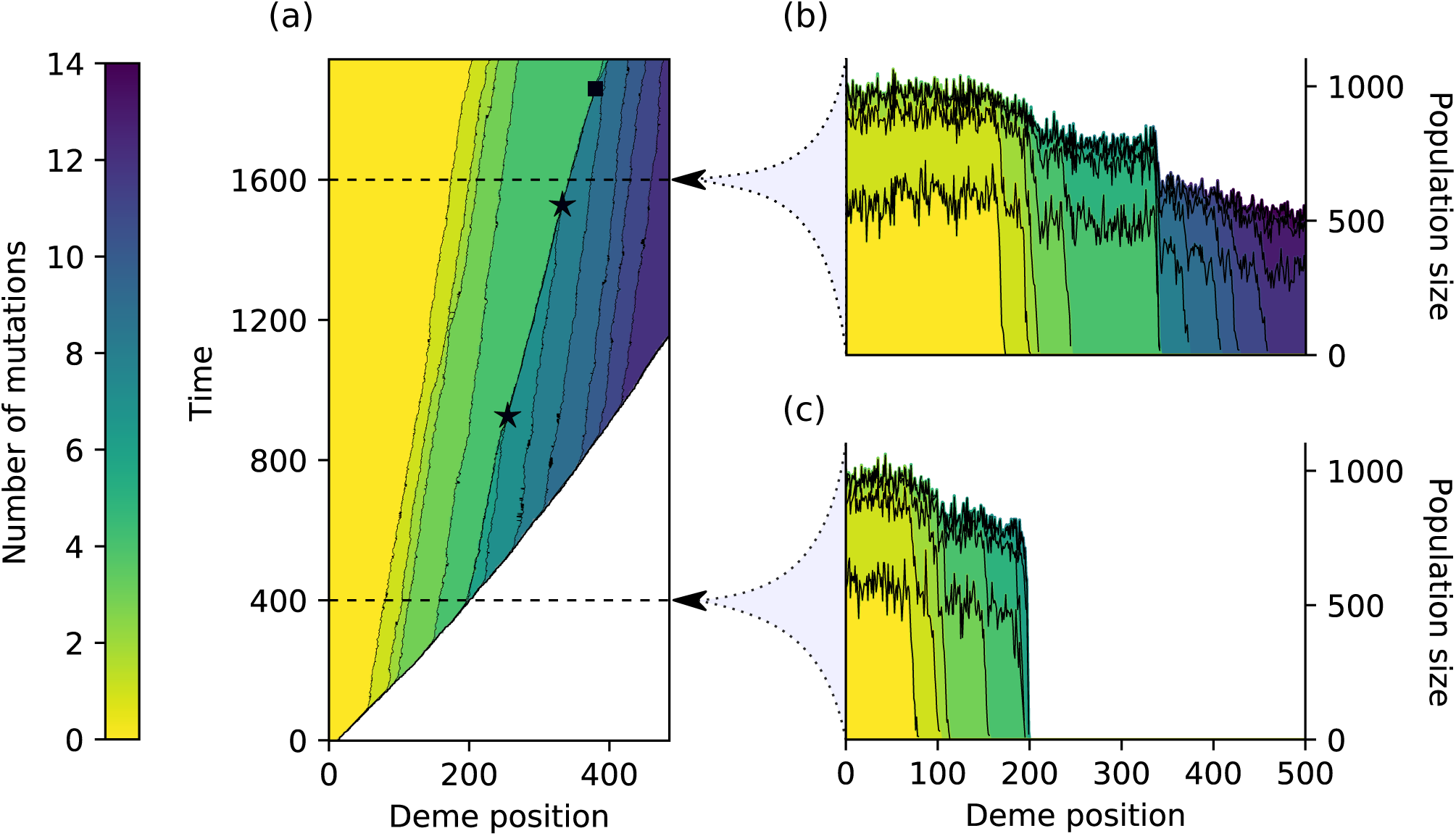
Typical simulation of the spatial Muller’s ratchet. (a) Time evolution of the population; in each row, the colour gives the number of mutations of the fittest individual in the deme, i.e., the number of clicks of the ratchet in this deme. Black stars indicate genetic waves collisions, and the black square indicates an inner click of the ratchet. (b, c) Genetic composition of the population at time *t* = 1600 and t = 400 respectively; the number of individuals carrying a given number of mutations is given by the height of the region with the corresponding colour. Parameter values are *N* = 1000, *r* = 1, m = 0.1, *s* = 0.05, *μ* = 0.025, *B* = 0.

**Figure 5:**
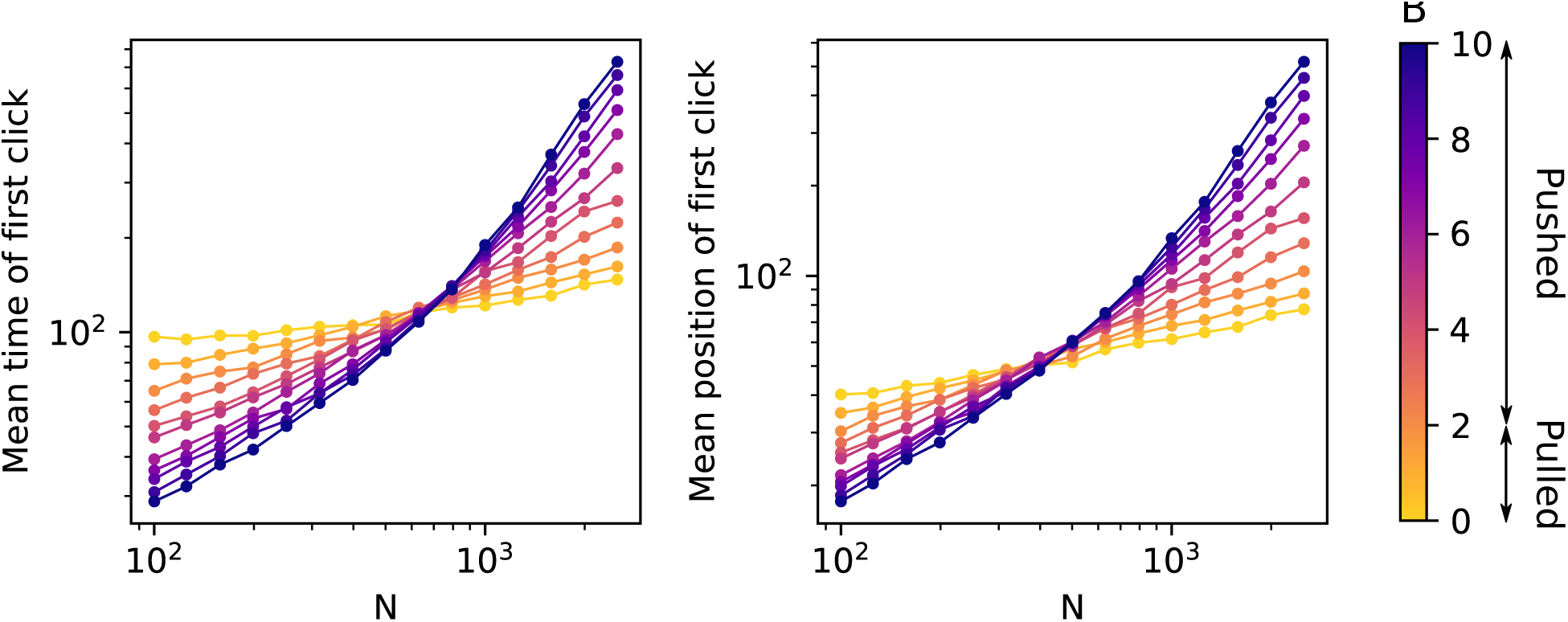
Scaling of the click rate and click position with *N*. The left plot shows the value of *T*_1_, and the right plot the value of 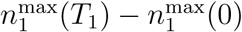, see Section 2 for the definitions. Each point is averaged over 5000 simulations. The parameter values are *r* = 1, *μ* = 0.01, *s* = 0.02, *m* = 0.1.

### Two-dimensional simulations

In the two-dimensional simulations, at *t* = 0 a five by five square of demes is occupied in the centre of the habitat and all other demes are empty. The number of individuals in these demes is chosen as in the one-dimensional case according to the deterministic values computed in (6). The simulation is run until *t* = 150. (Recall that *t* = 150 refers to the “actual time” of the simulation, and not to a number of iterations.) We want to record the number of clicks of the ratchet at the colonization time in each deme. One naive way of doing this could be to record for each deme the number of mutations of the first individual that migrates to this deme. However this would produce an extremely noisy picture. Even if the ratchet has not clicked yet, many individuals in the front carry mutations and could by chance migrate first to a new deme. In order to reduce this noise, we have chosen to look at the population at each time unit of the “actual time”, i.e., at *t* = 1, 2, …, 150, and to record the least number of mutations in each newly colonized deme. More precisely, let *n*_*i,j*;*k*_(*t*) denote the number of individuals carrying *k* mutations in deme (*i, j*), and let *N*_*i,j*_ be the total number of individuals in deme (*i, j*). We define the colonization time 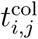 and number of clicks at colonization 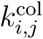 in deme (*i, j*) to be

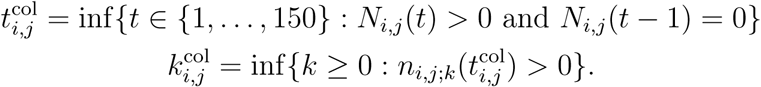

## 3 Results

### 3.1 Analysis of the deterministic limit

In this section we study the set of reaction-diffusion equations that we obtained by taking the deterministic scaling of the model, namely equation (2). Similar reaction-diffusion equations have been widely used to model biological invasions [43, 17]. They are usually studied through their travelling wave solutions. In our context, a travelling wave solution to (2) is a solution (*u*_*k*_)_*k*≥0_ that can be written

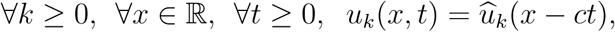

where *c* > 0 is the wave speed and

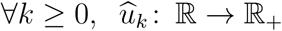

is the wave shape. A travelling wave solution is thus a constant wave form 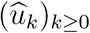 that is shifted at a constant speed *c* towards the positive reals. Additionally, we impose that the solution connects two stationary points of (2), i.e., that

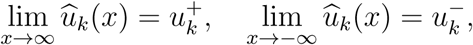

where 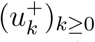 and 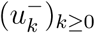 are two homogeneous solutions to (2). Let us first study the non-spatial equivalent of (2) to obtain the homogeneous solutions of the system.

#### Equilibrium of the non-spatial system

Let us consider the following non-spatial version of (2),

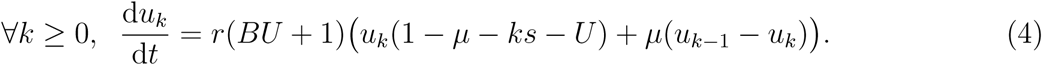

Equivalently, this system can be reformulated in terms of the total population size *U* and of the vector (*p*_*k*_)_*k*≥0_ = (*u*_*k*_*/U*)_*k*≥0_ giving the frequencies of the different types, that we call the *genetic composition* of the population. Equation (4) is then equivalent to

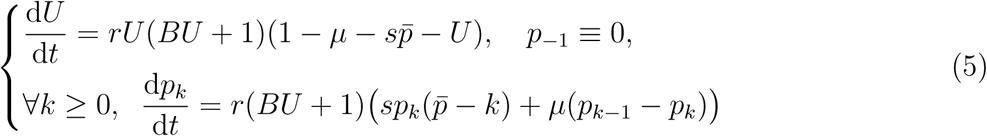

where we have set

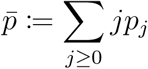

to be the mean number of mutations. Up to the non-constant population size, equation (5) has already been derived in [11] to describe dynamics of the frequencies of individuals carrying different numbers of mutations in Muller’s ratchet.

It is straightforward to see that if 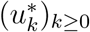 is a stationary point of (4), then either it is the trivial null equilibrium, or there exists some *k*_0_ ≥ 0 such that

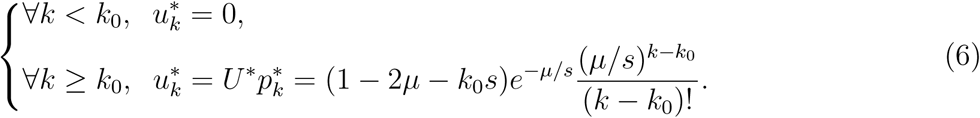

Thus, at the equilibrium, the population size is *U*^∗^ = 1 − 2*µ* − *k*_0_*s*, and the number of mutations has a Poisson distribution with parameter *µ/s*, shifted by *k*_0_, where *k*_0_ is the number of mutations of the best class.

Recall that a travelling wave solution should connect two stationary points of (4). From the above calculation, we conclude that equation (2) has at most two different types of travelling waves. Travelling waves that connect the trivial null equilibrium with a non-trivial equilibrium of the form (6). Travelling waves connecting two equilibria of the form (6) for different values of *k*_0_. The former travelling wave corresponds to the expansion of a population in an empty available habitat. We call it a *population travelling wave*. The latter wave corresponds to the invasion of fitter individuals in a region where the ratchet has clicked. The total population size remains almost constant, but the genetic composition of the population shifts from one Poisson equilibrium to the other. We call it a *genetic travelling wave*. Let us now study these two kinds of waves separately.

#### Population travelling wave

We first prove the existence of population travelling waves. We can write (2) in terms of the total population size and of the genetic composition. Equation (2) is then equivalent to

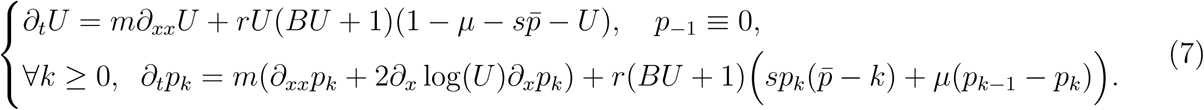

Suppose that the initial genetic composition is Poisson with parameter *µ/s* for all *x* ∈ ℝ. Then, as the Poisson distribution is a stationary point of (4), (*p*_*k*_)_*k*≥0_ remains Poisson for all *x, t*, and the equation for *U* now reads

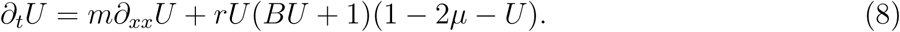

Up to a scaling in time and space, the above equation has already been considered in [5, 24], and we know from [24] that it admits a travelling wave solution for all speeds *c* ≥ *c*_0_, where *c*_0_ is given by

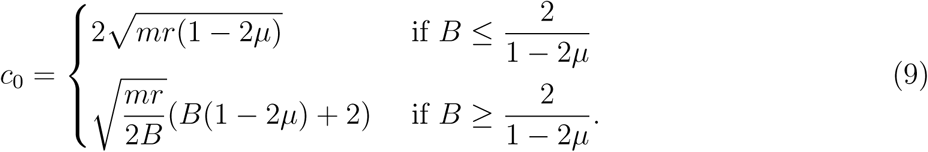

Thus, if 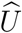 is the wave form of a travelling wave solution with speed *c* of (8), then

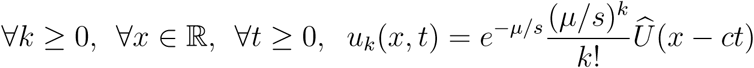

is a population travelling wave solution to (2). In this case, for *B* ≤ 2*/*(1 − 2*µ*), the population wave is pulled, as it has the same minimal speed as the linearized version of (2), while for *B* ≥ 2*/*(1 − 2*µ*) the wave is pushed. In addition to the existence of travelling wave solutions to (8), there exist several results concerning the convergence to these travelling waves for various initial conditions, see e.g. Theorem 4.1 and Theorem 4.3 from [1]. These results can be directly adapted to the solutions of (2), started from a Poisson genetic composition.

As a remark, the above population travelling wave connects the null equilibrium with the equilibrium (6) for *k*_0_ = 0. In a similar way we can find a population travelling wave for all *k*_0_ ≥ 0, the corresponding wave speed is obtained by replacing the term 2*µ* by 2*µ* + *k*_0_*s*.

#### Genetic travelling wave

We simulated numerically equation (2) with initial condition

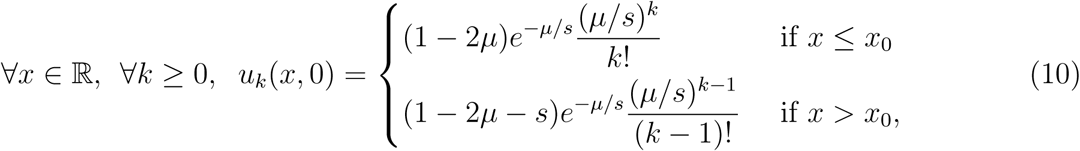

for some *x*_0_ ∈ ℝ, see Figure 3. The population is initially divided into two regions, one towards the positive reals where the fittest individuals carry one deleterious mutation, the other towards the negative reals where the fittest individuals carry no mutation. The initial condition of the former region approximates the state of the population after one click of the ratchet, while that of the latter region approximates the state of the population when no click has occured. Thus, equation (10) corresponds to the situation where fit individuals from the bulk invade a region where the ratchet has clicked once. A travelling wave rapidly forms at the onset of the simulation. Such a wave connects the equilibrium given by (6) with *k*_0_ = 0, to that with *k*_0_ = 1, and corresponds to a genetic travelling wave.

We do not prove the existence of such a wave. Nevertheless, we are able to give an upper bound on the speed at which individuals with no mutations spread. Let us suppose that the total population size *U* is non-increasing in space, as observed in the simulations. Then, the following bounds would hold,

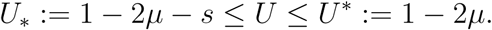

Using these bounds in the equation for *u*_0_ we obtain

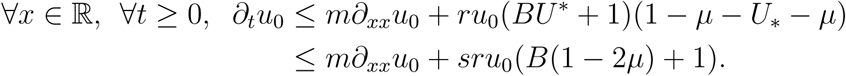

A classical comparison argument (see Proposition 2.1 in [1]) now shows that *u*_0_ is bounded above by the solution to the linear equation

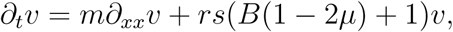

with initial condition 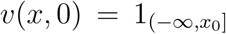. This linear equation can be solved explicitly, and an argument taken from [43], that we have recalled in Appendix B, shows that if *u*_0_ is spreading at speed *c*, then

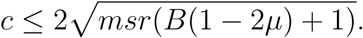

Comparing this bound with (9), we see that for a genetic travelling wave, 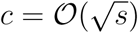, while for a population travelling wave, *c* = 𝒪(1). As we assume weak selection, i.e., *s* ≪ 1, genetic travelling waves are much slower than population travelling waves.

### 3.2 Simulations of the model

#### Spatial clicks at the front

In order to reproduce a range expansion, we considered a population initially at carrying capacity and mutation-selection balance, and then let it expand into an empty region of space. A typical simulation output is shown in Figure 4. At the start of the simulation, the population is invading the new habitat at a constant speed, forming a stochastic population travelling wave, see Figure 4, panel (a). Within each deme, the total population size and the genetic composition fluctuate around their deterministic values derived in (6), see Figure 4 panel (c).

Eventually, due to stronger genetic drift at the front, the best type is lost from the front. The population is now divided into two spatial regions: one region towards the front, that has lost the best type; the other region towards the bulk where the best type remains present. By analogy with the non-spatial Muller’s ratchet, we call this loss of the best type at the front a spatial click of the ratchet. The region where the ratchet has clicked rapidly approaches a Poisson distribution of mutations, with a slightly decreased total population size as predicted by (6).

The situation is now a mixture of the two initial conditions considered in Figure 3. The population has not yet colonized all the available demes, and it keeps on spreading according to a population travelling wave (whose speed is decreased due to the spatial click). Nevertheless, the population is now divided into two regions in a similar way to Figure 3 panel (b), and we see the formation of an inner genetic wave. The genetic wave is much slower than the population wave.

This can be understood from the calculations of Section 3.1. We have shown that the speed of genetic waves scales as 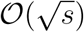, and thus vanishes as the selection coefficient goes to 0. Conversely, for a fixed ratio *µ/s*, the minimal speed of population waves, provided in equation (9), converges to a positive limit as *s* goes to 0. Thus, as we consider *s* ≪ 1, we expect genetic waves to be much slower than population waves, as observed.

After the first spatial click of the ratchet, the population returns to its original selection-mutation balance, except that each individual now bears an additional mutation. Eventually a new spatial click of the ratchet will occur, and subsequent clicks will occur repeatedly during the expansion, see Figure 4. We can interpret these results in terms of expansion load. Recall that the expansion load refers to the additional loss in fitness due to range expansion. Initially, the population is at mutation-selection balance, and has a mutational load *µ/s*. After *k* clicks of the ratchet, the mean number of mutations at the front is *µ/s* + *k*. Thus in our context, the expansion load is given by the number of spatial clicks that the population has experienced. In order to quantify the speed at which expansion load is building, we need to compute the rate at which spatial clicks of the ratchet happen in the population.

#### Bulk dynamics

After the ratchet has clicked several times, the bulk is divided into regions where the number of mutations of the fittest individuals corresponds to the number of clicks that the region has experienced. Each region has a fitness advantage compared to the adjacent region located towards the front, but has a fitness disadvantage compared to that towards the bulk. Thus each region is able to move forward even as it is being chased, resulting in a sequence of genetic travelling waves, see Figure 4 panel (a). We are interested in the dynamics of these genetic waves.

Each wave separates two regions that have accumulated distinct numbers of mutations, and thus have distinct mean fitness. The speed of the wave increases with the fitness difference between these regions. After a single click of the ratchet, this fitness difference is *s*. All waves that separate regions where the ratchet has clicked once spread at the same average speed, leading to the parallel genetic waves observed in Figure 4 panel (a). However, we observe that “double clicks” of the ratchet occur: the best and second best class of individuals can be lost simultaneoulsy from the front, for example this is the case at *t* = 400 in Figure 4. In this case, the fitness difference between the two sectors resulting from the click is 2*s*, and the corresponding genetic wave spreads faster. It is able to catch up the next genetic waves, leading to wave collisions as indicated by the black stars in Figure 4. When two waves collide, the fitness difference between the two regions separated by the resulting wave increases, and thus the wave speeds up.

Interestingly, after several wave collisions, we observe that a genetic wave can split into two waves, as is indicated by the black square in Figure 4 (see also Figure 8 for an example of simulation where this split occurs earlier). This corresponds to an “inner” click of the ratchet: the best class of the genetic wave is lost from the front (of the genetic wave). The mechanism leading to such an inner click is the same as that leading to spatial clicks at the front of the population wave. Fit individuals at the front of a genetic wave have a high growth rate as they compete with individuals that have accumulated many deleterious mutations. If an individual from the second best class gets ahead, it can rapidly grow a large subpopulation that will further expand, creating a new genetic wave. We expect such an event to occur at a higher rate when the fitness difference between the sectors separated by the genetic wave is large.

**Figure 6:**
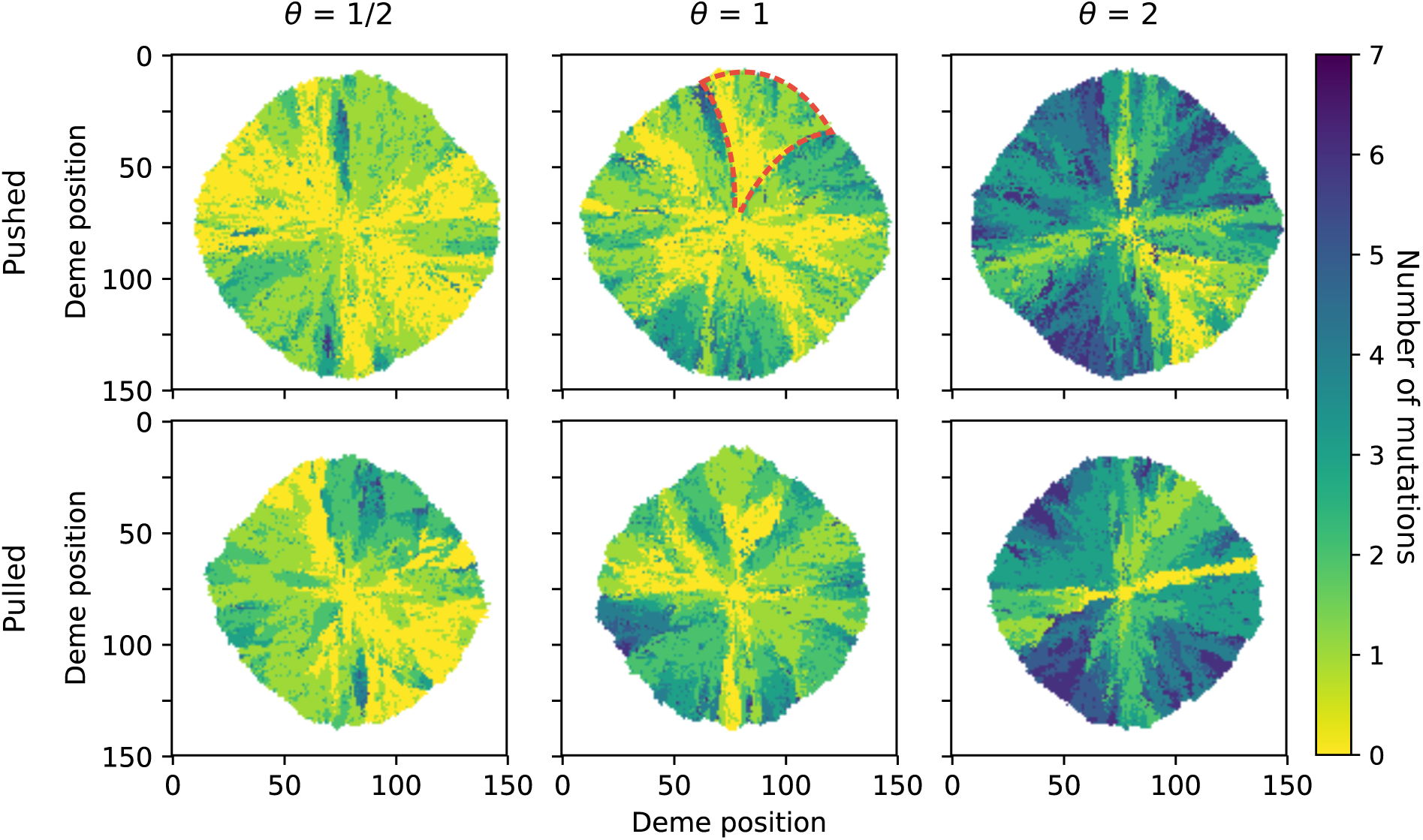
Two dimensional simulations, shown at time *t* = 150. All simulations are realized with *N* = 300; *μ* = 0.02; *r* = 1, *m* = 0.1. We set *B* = 0 in the pulled case, and B = 3 in the pushed case. The value of the selection coefficient is chosen in *s* ∈ {0.01, 0.02, 0.04} to obtain the various values of *μ/s*. The colour indicates the number of mutations of the best type at the deme colonization, see Section 2. The red dashed line indicates a funnel-shaped sector.

**Figure 7:**
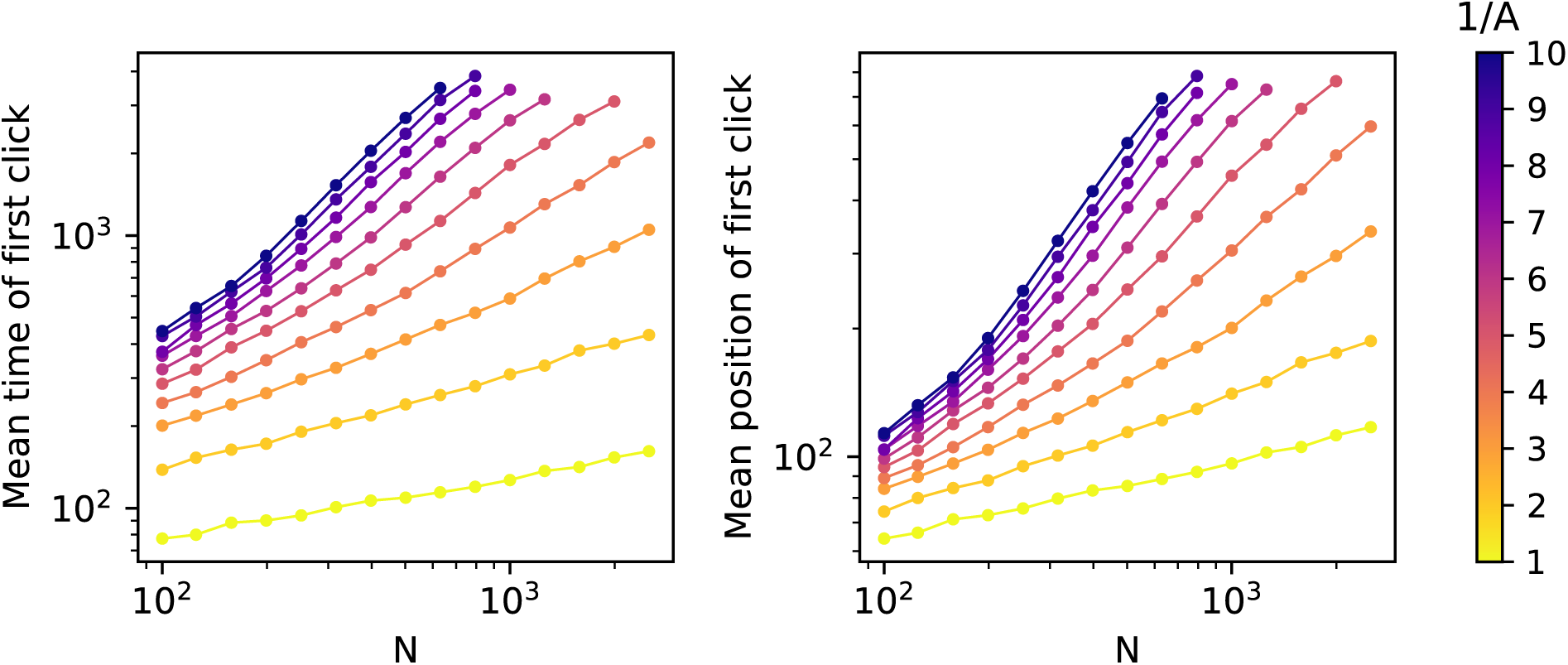
Scaling of the click rate and click position with *N* for the parametrization (11). The left plot shows the value of *T*_1_, and the right plot the value of 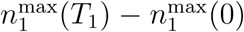, see Section 2 for the definitions. Each point is averaged over 5000 simulations. The parameter values are *ρ* = 1, *μ* = 0.01, *s* = 0:02, *m* = 0.1.

**Figure 8:**
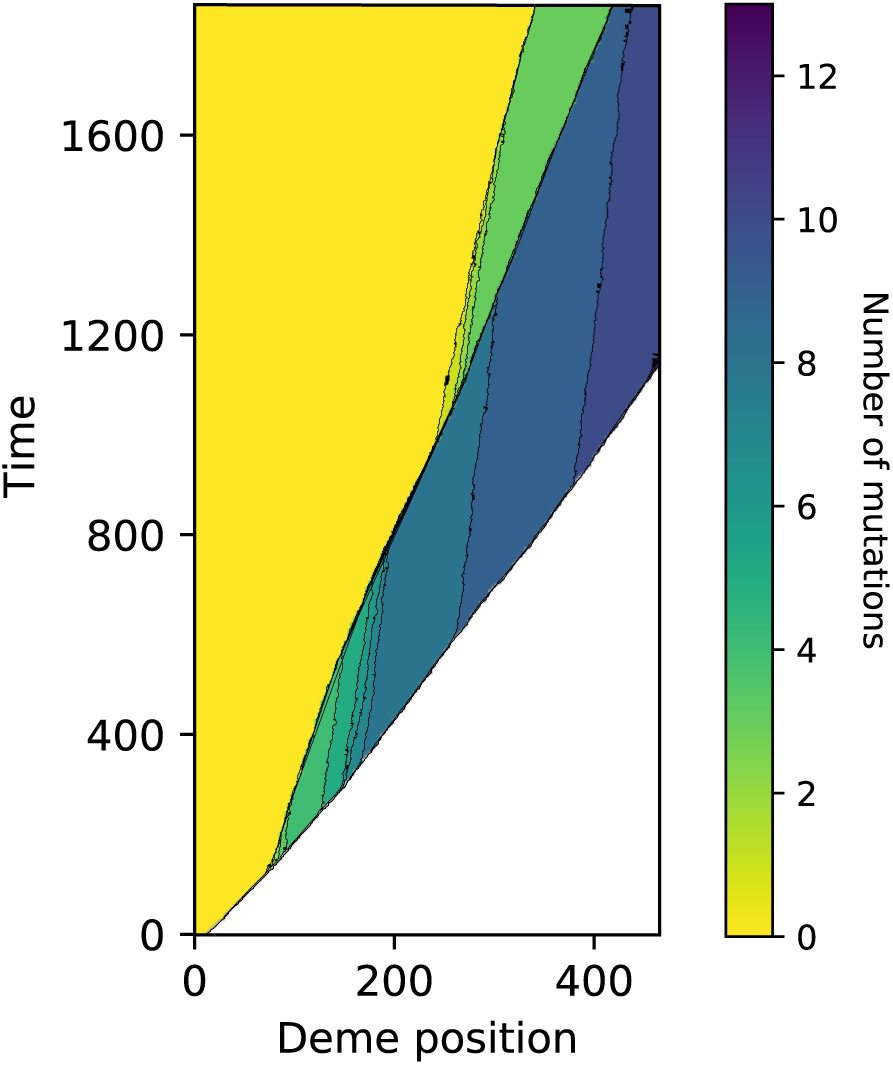
Space-time diagram of a simulation of the spatial Muller’s ratchet. Each row corresponds to a time value, and the number of mutations of the fittest individual in each deme at that time is represented. Notice that a fast genetic wave rapidly forms, and that it experiences several successive inner clicks of the ratchet. The parameter values are the same as in Figure 4.

The picture that emerges from the analysis of the dynamics of the bulk is the following. We can think of the bulk as a branching-coalescing system of particles: each genetic wave corresponds to a particle located at the front of that wave. Then the system has the following dynamics. Each particle follows a random motion, with an average speed towards the positive reals that increases with the fitness difference of the regions it separates. When two particles meet, i.e., when two genetic waves collide, they merge, and the resulting particle speeds up. Finally, when an inner click of the ratchet occurs, a new particle is created, and the two daughter particles are slower than their mother. Such a branching event occurs at a rate that increases with the fitness difference of the regions separated by the particle.

### 3.3 Impact of the Allee effect on the expansion load

We now aim to study the impact of an Allee effect on the expansion load, i.e., on the click rate in our context. Analysis of the formation of expansion load requires us to take into account genetic drift. Recall that a generator calculation suggests that a good approximation of our model that retains finite size fluctuations is given by equation (3). The parameter that controls the strength of the Allee effect is *B*. However, we see from equation (3) that increasing *B* also increases the noise term, and hence increases genetic drift. Thus the parameter *B* has two antagonistic effects on the click rate: on the one hand it should reduce the click rate by increasing the strength of the Allee effect, and shifting the nature of the wave from pulled to pushed; on the other hand, it increases the click rate by reinforcing genetic drift, and hence gene surfing. In order to disantangle these two effects, we will study the impact of *B* on the scaling with *N* of the click rate. If the pulled/pushed nature of the wave does not impact expansion load, increasing *B* should only increase the strength of the drift and we expect a similar scaling of the click rate with *N* for different values of *B*.

A direct computation of the click rate from (3) is not feasible. We thus used simulations to assess the impact of *B* on the click rate. Starting from an initial condition similar to Figure 4, we let the population expand, and record the time *T*_1_ and spatial location 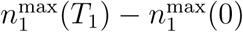 of the first spatial click of the ratchet, see Section 2 for a precise definition of these quantities. Both *T*_1_ and 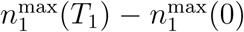 should be inversely related to the click rate. Figure 5 panel (a) shows the time of this first click, averaged over 5000 simulation replicates, for various values of *B* and *N*. We have performed a similar analysis for the second and third clicks of the ratchet. The results, shown in Figure 9, are qualitatively similar.

**Figure 9:**
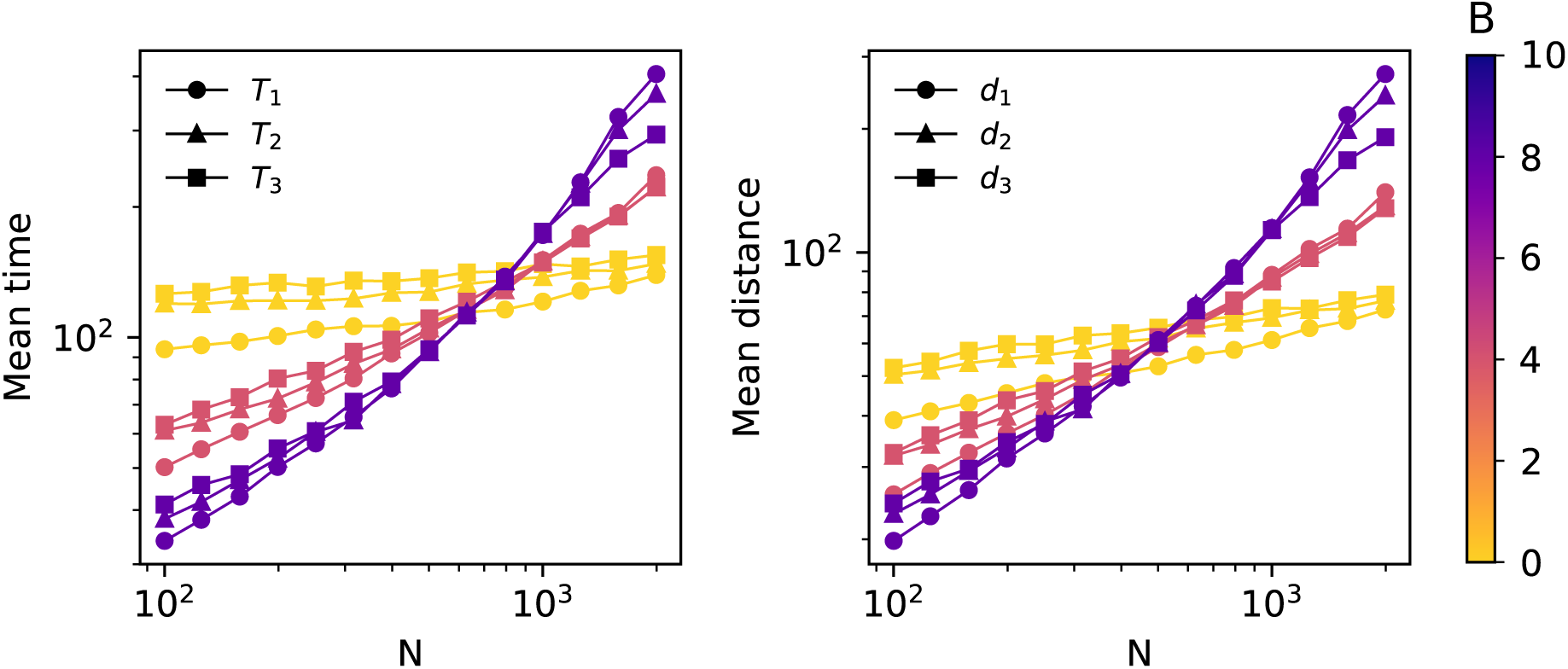
Scaling of the click rate and click distance with *N*. Each point is averaged over 5000 simulations. The parameter values are *r* = 1, *µ* = 0.01, *s* = 0.02, *m* = 0.1.

First, notice that for each fixed value of *B*, the mean time to the first click increases with *N*, i.e., the ratchet is slower for large population sizes. This is in agreement with our intuitive understanding of the ratchet, since increasing *N* reduces the strength of genetic drift and hence reduces gene surfing. Second, for a fixed value of *N*, increasing *B* either speeds up the ratchet if *N* is low, or slows it down if *N* is large. This observation can be explained intuitively as follows. Recall that *B* has two antagonistic effects on the click rate: increasing the genetic drift, and increasing the gene flow from the bulk to the front. For low values of *N*, the population size in the bulk is low, and the gene flow from the bulk restores the genetic diversity at the front less efficiently than for large values of *N*. Thus increasing the gene flow from the bulk to the front has a larger impact on the click rate for large values of *N*. For low values of *N*, the increase of genetic drift with *B* prevails, while the converse holds for large values of *N*.

Let us now consider the scaling of the time to the first click, *T*_1_, with *N*. First, notice that for any value of *B, T*_1_ scales faster than a power law with *N*. It is clear that *N* has more impact on *T*_1_ for pushed waves, i.e., for large *B*, than for pulled waves. In the pulled case, *T*_1_ increases with *N* very slowly, and the rate of ratchet is only slightly changed by the population size. Conversely, in the pushed case, *N* has a drastic effect on *T*_1_: the ratchet clicks very fast for low *N*, but we see almost no click of the ratchet for large *N*.

We thus conclude that the Allee effect has a large impact on the click rate. For small values of *B*, the population size has little impact on the building of the expansion load. This reflects the fact that the dynamics is mostly determined by the few individuals in the front, that are almost insensitive to the change in the carrying capacity in the bulk. Conversely, for large values of *B*, the dynamics of the wave is determined by an intermediate region between the bulk and the front. Increasing *N* reduces the genetic drift in this region and leads to the large effect of *N* on *T*_1_ observed in Figure 5.

## 4 Discussion

Expansion load originates from the strong genetic drift induced by the low population size at the edge of an expanding population. From a modelling perspective, demography, spatial structure, stochasticity and selection are minimal ingredients to account for expansion load. Each of these features is known to make mathematical treatment harder, and thus building a tractable model for expansion load is challenging. In this work we proposed a model similar in spirit to [40], but with two major differences: we greatly simplified the genetic structure of the population to that of a Muller’s ratchet, and we introduced an Allee effect in the population, tuned by the parameter *B*. This simplification allowed us to prove rigorous results for the deterministic scaling of the model, however an analysis of the stochastic scaling (3) where the building of an expansion load occurs remained out of reach.

Among other factors that are known to impact the genetics of range expansion, such as density dependent migration [6] or long distance dispersal [15], we have focused here on the impact of an Allee effect on expansion load. It is already understood from several studies that an Allee effect impedes gene surfing [5, 43, 18]. In agreement with these findings and our intuitive expectations, we have shown that adding an Allee effect to the population slows down the rate at which the ratchet clicks for large population size, and thus reduces the expansion load.

However, [43, 18] predict a sharp qualitative difference between pulled and pushed waves. This disagrees with Figure 5 where increasing *B* continuously changes the scaling of the click rate with *N*. Nevertheless, note that the results in [43, 18] were obtained in a deterministic setting, and that stochasticity has a tendency to smoothen such transitions. In a stochastic setting, [5] have fitted a power law to the rate of genetic diversity loss in an expanding population, as a function of *N*. They predicted that the exponent of this power law remains constant outside of the parameter region *B* ∈ [2, 4], see their Figure 4 and Figure 5. Again, in our Figure 5 we see a change in the scaling on the entire range of *B*. This discrepancy might be explained by the coupled effect *B* has on the nature of the wave and the genetic drift.

Moreover, for low *N*, we observe that increasing *B* increases the rate of the ratchet. This originates from the complex interaction between Allee effect and genetic drift in our model. More generally, genetic drift depends on the rate at which birth and death events occur in the population. Changing the strength of the Allee effect should modify these rates, and we can expect the Allee effect to interact with genetic drift for a large class of models. The specific form of this interaction should depend on the details of the microscopic model under consideration and the way the Allee effect is implemented. Therefore, we believe that our results cannot be directly transposed to other models of population expansion incorporating an Allee effect. The impact of the Allee effect on the expansion load should depend in a crucial way on its interplay with genetic drift, which is a model dependent feature. In order to illustrate this, we have reproduced the results of Figure 5 for an alternative parametrization of our model, see Section C and Figure 7. We observe that the results are qualitatively very different. Increasing the strength of the Allee effect reduces the rate of the ratchet for all *N* in this new parametrization.

Our model also relates to the vast literature on Muller’s ratchet, see e.g. [35] and references therein. The effect of spatial structure on the dynamics of Muller’s ratchet has already been investigated in [30]. They concluded that, for a fixed total population size, subdividing the population into smaller habitats reinforces Muller’s ratchet, since genetic drift is enhanced in each single habitat. The setting we consider is different. The total population size is not fixed as we allow new subpopulations to grow in empty demes. We find that space has two effects on the ratchet. In the bulk of the population, we do not observe clicks of the ratchet. Spatial structure has a stabilizing effect: if the ratchet clicks in one deme, the best type can be reintroduced by migration from the adjacent demes. Conversely, in the front spatial structure causes low population size and thus speeds up the ratchet. Overall, our study shows that spatial structure can interact in a non-trivial way with Muller’s ratchet.

We have considered the expansion of a population in a linear one-dimensional habitat. Multiple studies have also been concerned with range expansion in two dimensions, using microbial growth experiments on petri dishes [27, 28, 33, 23] or simulations [13, 39]. The typical set up of these studies is to place a drop containing two labelled strains in the centre of a petri dish, and to let them expand. The colony is rapidly separated into sectors where only one of the two strains is present, and the other is absent, see for example Figure 1 in [23]. These studies have examined the dynamics of these sectors, especially when there exists a fitness difference between the two strains. They have established that the boundary between two sectors should move towards the strain with the lower fitness, and gave an expression for the speed of the boundary in terms of the fitness difference [33, 27].

In the context of the spatial Muller’s ratchet, the major expected difference between one and two dimensions is the following. In one dimension, once the ratchet has clicked, best type individuals are trapped in the bulk of the population. The only way to restore the fitness at the front is that the genetic wave of fit individuals catches up with the population wave. We know that this is extremely unlikely, because the genetic wave is much slower than the population wave. In two dimensions, the front is a one-dimensional curve, and a click of the ratchet only removes best type individuals from a small part of it. The remaining best type individuals have a fitness advantage compared to individuals in demes where the ratchet has clicked, and according to the aforementioned studies, they should be able to remove the unfit individuals from the front. Thus, in two dimensions, a click of the ratchet does not irredeemably trap best type individuals in the bulk, and fitness should be restored by migration of fit individuals from parts of the front where the ratchet has not clicked.

In order to assess these predictions and to compare the behavior of our model to previous studies, we have simulated the spatial Muller’s ratchet on a two dimensional lattice. Demes are now indexed by ℤ^2^, the reproduction rules within each deme remain the same, but at each migration event individuals now choose one of the four adjacent demes with equal probability. A key difference between our model and the microbial growth experiments is that the cells are non-motile and unable to migrate. In the spatial ratchet, the bulk of the population is dynamical, and fit individuals slowly expand according to genetic waves. In the growth experiments, cells in the bulk remain at their initial location, and the observed patterns correspond to a “frozen record” of the front at the time of colonization. In order to carry out the comparison between our model and the existing studies, we have depicted in Figure 6 the number of mutations in each deme at its colonization time, see Section 2 for the precise definition of this quantity. We emphasize the fact that this is *not* the state of the entire population at the end of the simulation: many sectors will have been taken over by fit individuals from the bulk and their shapes would not be comparable to that of the sectors obtained in microbial growth experiments. For comparison, we have shown the state of the population in Figure 10.

**Figure 10:**
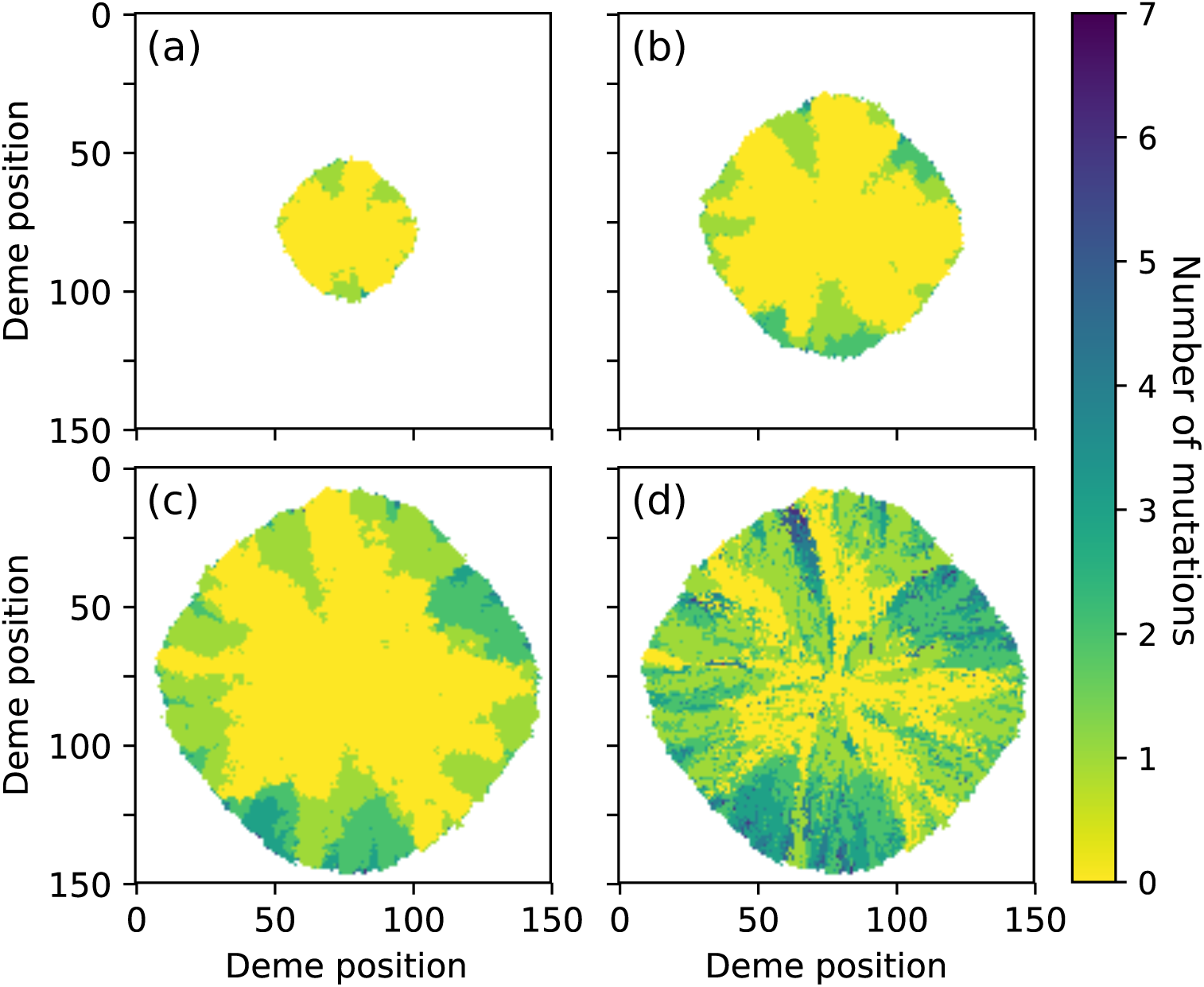
Simulation of the two-dimensional spatial Muller’s ratchet. (a,b,c) State of the population at time *t* = 50, 100, 150 respectively. In each deme the number of mutations of the fittest individuals is represented. (d) Number of mutations at colonization time, see Section 2 for the definition. The parameters are *N* = 300, *µ* = 0.02, *s* = 0.02, *r* = 1, *m* = 0.1, *B* = 0.

As in the one-dimensional case, we observe in Figure 6 clicks of the ratchet leading to the formation of sectors with lower mean fitness. An achievement of the microbial growth experiments on petri dishes is to link the shape of these sectors to their relative fitness. If the strains that are placed on the petri dish have the same fitness, then the sectors should be “cone-shaped”: the boundary between two sectors is wandering due to stochastic effects but does not have a preferential direction. Conversely, if one strain is fitter than the other, the sectors of the fitter strain should have a typical “funnel” shape, see for instance Figure 4 in [23] for examples of these two shapes. Most sectors in Figure 6 have a cone shape, indicating that the expansion is nearly neutral. Selection is too weak in these simulations to allow fit individuals to efficiently remove unfit individuals from the front. Nevertheless, we have indicated by a red dashed line a sector that has the typical shape of a selectively advantaged strain. Notice that the regions adjacent to this sector have experienced multiple clicks of the ratchet, and thus that the fitness advantage of this sector is large.

Apparently, in the parameter region we have considered, selection is not strong enough to reverse spatial clicks of the ratchet and efficiently restore fitness at the front. The dynamics of the sectors is nearly neutral, and we do not observe any major difference from the one-dimensional case. The pushed/pulled nature of the wave seems to have the same qualitative effect on the clicks of the ratchet as in the one-dimensional case. A better understanding of the two-dimensional case would require a more quantitative and thorough investigation, which goes beyond the scope of the present work.

During a range expansion, the front can accumulate mutations leading to an expansion load, but individuals in the bulk do not bear this additional burden. Thus, at each location in space, fitness should be slowly recovered through migration of fit individuals from the bulk: expansion load is a transient phenomenon [20, 39]. In the spatial Muller’s ratchet we have a clear quantification of the rate of this fitness recovery. Fit individuals take over the population through inner genetic waves, with a speed proportional to the square root of their selective advantage. As discussed previously, on the one hand the speed of genetic waves can increase due to wave collisions. On the other hand, the speed of the population wave is decreased by the successive clicks of the ratchet. It is natural to ask whether the population wave is eventually caught up by a genetic wave. More generally, it would be interesting to study the long-term behavior of the spatial Muller’s ratchet. A possible starting point is to approximate the dynamics of the bulk of the population by a branching-coalescing particle system as described previously, and to study the asymptotic behavior of this simplified system.

The nature of population travelling waves changes from pulled to pushed at the critical value of *B* = 2*/*(1 − 2*µ*). It is interesting to ask whether such a transition occurs for genetic waves. From (7), the per-capita growth rate of *p*_0_, the frequency of individuals without mutations, is

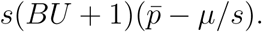

In a genetic travelling wave, the total population size *U* is almost constant (it ranges from 1 − 2*µ* to 1 − 2*µ* − *s*). As we have the constraint ∑*p*_*i*_ = 1, we see that, roughly speaking, the maximal per-capita growth rate of *p*_0_ is achieved for lower values *p*_0_. In our intuitive definition of pulled and pushed waves, this corresponds to the pulled case. In our model, genetic waves are always pulled, regardless of the value of *B*.

This essentially comes from the fact that we have considered a haploid population. The dynamics of the frequency *p* of a gene with fitness advantage *s* in a haploid population with local migration is given by the classical Fisher-KPP [17, 32] equation

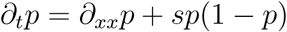

which is the archetypical example of a pulled wave. Considering a diploid population where homozygotes have fitness 1 + 2*αs* and heterozygotes a fitness 1 + (*α* − 1)*s* leads to a special case of the so-called Allen-Cahn equation [2]

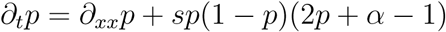

that displays a transition from pulled to pushed waves when varying the parameter *α*. One could thus obtain pushed genetic waves by considering a diploid population with heterozygote advantage [3].

## 5 Acknowledgment

We greatly thank Julien Berestycki for helpful discussions, and the editor and an anonymous referee for helpful comments. We are also grateful to the INRA MIGALE bioinformatics facility (MIGALE, INRA, 2018. Migale bioinformatics Facility, doi: 10.15454/1.5572390655343293E12) for providing computational resources.

## A Generator computations

### A.1 Deterministic scaling

Let (*n*_*i,k*_(*t*); *t* ∈ ℝ_+_, *i* ∈ ℤ, *k* ≥ 0) be the process described in Section 2, and recall that we have defined the renormalized process as

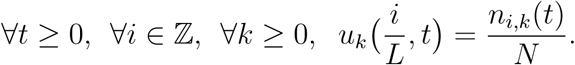

Let *ϕ* : ℝ → ℝ be a twice-differentiable function with compact support, and set

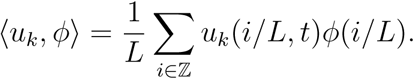

Recall that *U* stands for the total renormalized population size

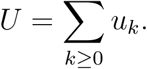

We expect a deterministic limit as *N* → ∞, so in order to identify the limit of the *u*_*k*_ under this scaling, it is enough to consider the generator applied to functions of this form. If *G*_*N*_ is the generator of *u*_*k*_, then

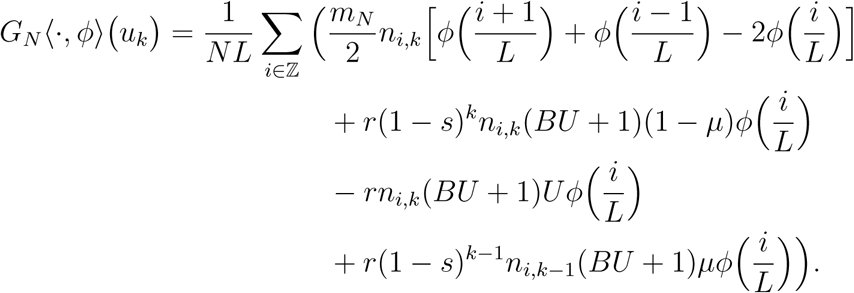

Thus we see that, provided *u*_*k*_ is converging and *m*_*N*_ */L*^2^ → *m*, the above quantity converges to 

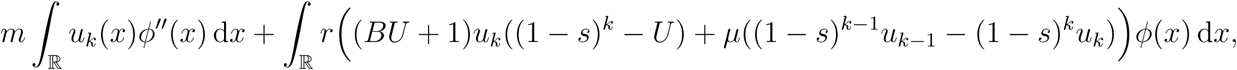

which suggests that in the limit (*u*_*k*_) solves

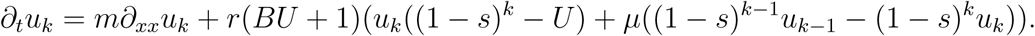

### A.2 Stochastic scaling

Consider a function *ϕ* : ℝ^ℤ × ℕ^ → ℝ that only depends on a finite number of coordinates, and suppose that *ϕ* is twice continuously differentiable. Under these assumptions, by making a Taylor expansion of *ϕ* at the point **u** = (*u*_*i,k*_; *i* ∈ ℤ, *k* ≥ 0) and ignoring terms of order greater than 1*/N* ^2^, we obtain the following expression for the generator 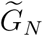 of the process with population size *N*

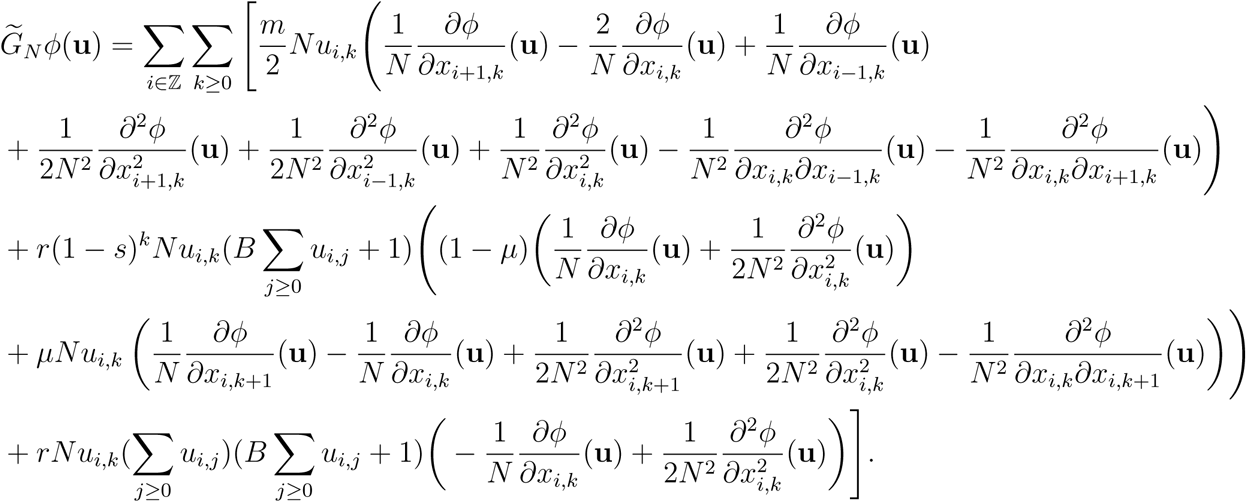

We suppose *m* ≪ *r* and that *µ* ≪ 1, so that we can neglect the mixed second order derivatives and discarding terms of 𝒪(1*/N* ^2^) we find that the generator of our rescaled process is approximately

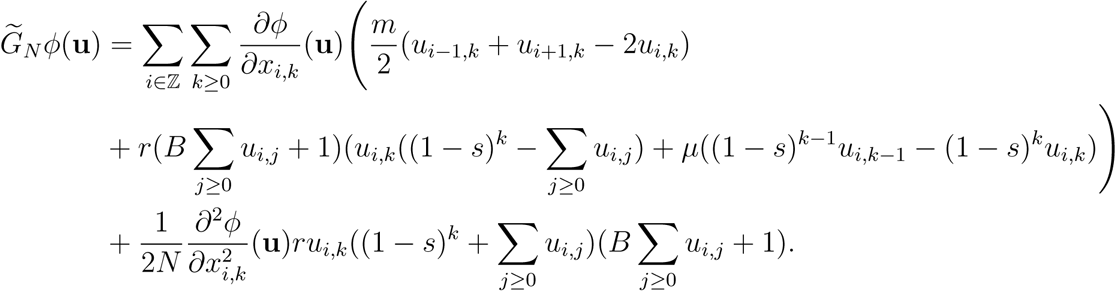

Thus, for large *N*, and *m* fixed such that *s, µ* ≪ 1, *m* ≪ *r*, our process is well-approximated by the following set of stochastic differential equations,

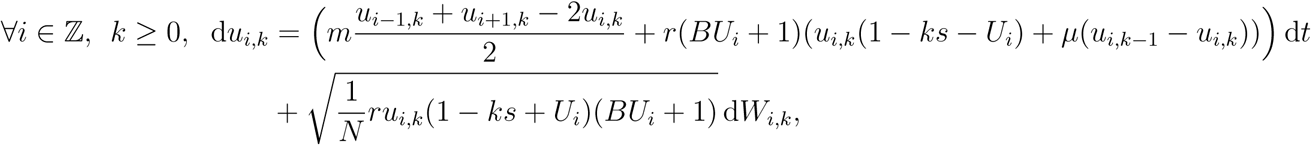

where (*W*_*i,k*_; *i* ∈ ℤ, *k* ≥ 0) are independent Brownian motions.

Writing

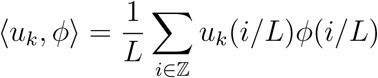

as before and speeding up time by a factor of *L*^2^ in order to obtain a diffusive rescaling and scaling *N* ↦ *LN* (corresponding to replacing the population size in a deme by the local population density), we expect that as *L* → ∞ we should recover the system of stochastic partial differential equations

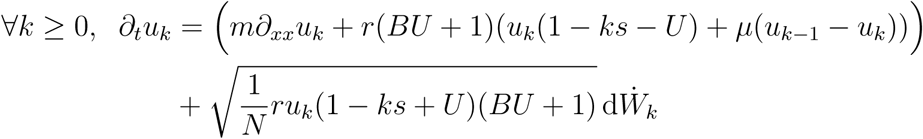

where 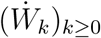 are independent space-time white-noises, see for example [4], Section 2. We emphasize that this derivation is heuristic. Again, see [9, 36] for rigorous treatments of similar convergence results.

## B Spread of a linear wave

Let us consider the equation

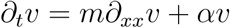

with bounded initial condition *v*(*x*, 0) = *v*_0_(*x*). The solution to this equation is

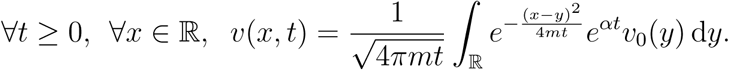

Following [43], let *c* ≥ 0 be some speed, then

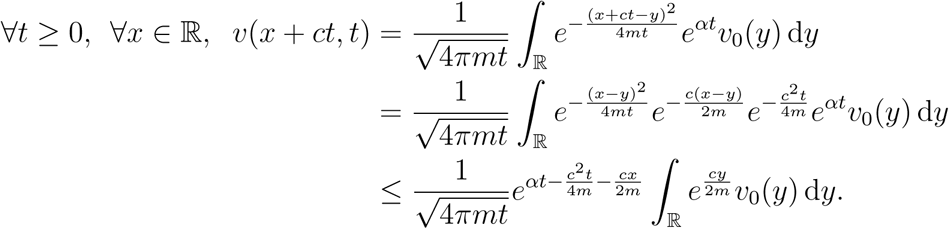

Thus, for 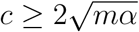,

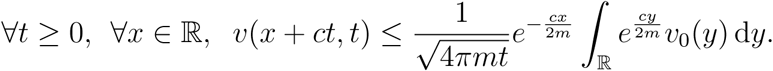

Hence, provided the integral is finite and 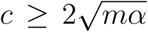 (which is the case when *v*_0_ is Heaviside), *v*(*x* + *ct, t*) goes to 0 uniformly on sets of the form [*A*, ∞] for *A* ∈ ℝ. This shows that the process *u* of Section 3.1 cannot converge to a travelling wave solution with speed larger than 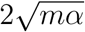, i.e., larger than 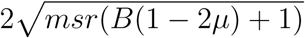 in this case.

## C Alternative parametrization of the Allee effect

In the current work, the birth and death rates have been chosen so that the overall growth rate of the population is a cubic function of the population size *n*,

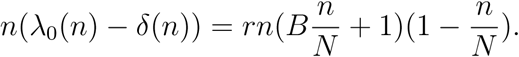

An alternative parametrization of the same polynomial can be obtained by setting *ρ* = *rB* and *A* = 1*/B*, so that

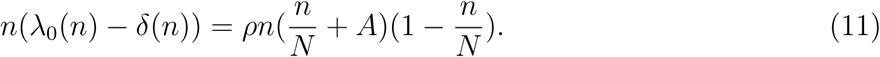

In this case, the population exhibits a weak Allee effect for *A* ∈ (0, 1) and no Allee effect for *A* ≥ 1, so that the strength of the Allee effect is inversely related to the parameter *A*.

We have reproduced the results of Figure 5 for this alternative parametrization. The results are shown in Figure 7. In order to make the comparison with Figure 5 easier, we have used the values of *A* corresponding to the values of *B* used in Figure 5. The result is qualitatively very different from Figure 5. Increasing the strength of the Allee effect reduces the click rate for all *N*, whereas this was only the case for large *N* in Figure 5.

## Supplementary figures

### Successive clicks of the ratchet

Recall the expression for the time *T*_1_ to the first click given in Section 2. Similarly, we can define the time between the *k* − 1-th and the *k*-th click as

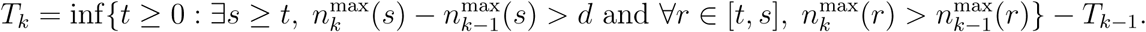

The number of demes that the population has colonized between the *k* − 1-th and the *k*-th click is then given by

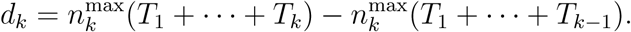

Figure 9 shows the values of *T*_1_, *T*_2_, *T*_3_, and *d*_1_, *d*_2_, *d*_3_, averaged over several simulations, for different values of the parameters *B* and *N*.

## References

[1] Donald G. Aronson and Hans F. Weinberger. Nonlinear diffusion in population genetics, combustion, and nerve pulse propagation. In Partial Differential Equations and Related Topics, pages 5–49. Springer Berlin Heidelberg, 1975. doi: 10.1007/BFb0070595.

[2] Nicholas H. Barton. The dynamics of hybrid zones. Heredity, 43:341–359, 1979. doi: 10.1038/hdy.1979.87.

[3] Nicholas H. Barton and Michael Turelli. Spatial waves of advance with bistable dynamics: Cytoplasmic and genetic analogues of allee effects. The American Naturalist, 178:E48–E75, 2011. doi: 10.1086/661246.

[4] Nick H. Barton, Frantz Depaulis, and Alison M. Etheridge. Neutral evolution in spatially continuous populations. Theoretical Population Biology, 61:31–48, 2002. doi: 10.1006/tpbi.2001.1557.

[5] Gabriel Birzu, Oskar Hallatschek, and Kirill S. Korolev. Fluctuations uncover a distinct class of traveling waves. Proceedings of the National Academy of Sciences, 115:E3645–E3654, 2018. doi: 10.1073/pnas.1715737115.

[6] Gabriel Birzu, Sakib Matin, Oskar Hallatschek, and Kirill S. Korolev. Genetic drift in range expansions is very sensitive to density feedback in dispersal and growth. 1903.11627 2019. URL https://arxiv.org/pdf/1903.11627.

[7] David S. Boukal and Ludĕk Berec. Single-species models of the Allee effect: Extinction boundaries, sex ratios and mate encounters. Journal of Theoretical Biology, 218:375–394, 2002. doi: 10.1006/jtbi.2002.3084.

[8] Ron Do, Daniel Balick, Heng Li, Ivan Adzhubei, Shamil Sunyaev, and David Reich. No evidence that selection has been less effective at removing deleterious mutations in Europeans than in Africans. Nature Genetics, 47:126–131, 2015. doi: 10.1038/ng.3186.

[9] Rick Durrett and Wai-Tong (Louis) Fan. Genealogies in expanding populations. The Annals of Applied Probability, 26:3456–3490, 2016. doi: 10.1214/16-AAP1181.

[10] Christopher A. Edmonds, Anita S. Lillie, and L. Luca Cavalli-Sforza. Mutations arising in the wave front of an expanding population. Proceedings of the National Academy of Sciences, 101:975–979, 2004. doi: 10.1073/pnas.0308064100.

[11] Alison M. Etheridge, Peter Pfaffelhuber, and Anton Wakolbinger. How often does the ratchet click? Facts, heuristics, asymptotics, page 365–390. London Mathematical Society Lecture Note Series. Cambridge University Press, 2009. doi: 10.1017/CBO9781139107020.016.

[12] Laurent Excoffier and Nicolas Ray. Surfing during population expansions promotes genetic revolutions and structuration. Trends in Ecology & Evolution, 23:347–351, 2008. doi: 10.1016/j.tree.2008.04.004.

[13] Laurent Excoffier, Matthieu Foll, and Rémy J. Petit. Genetic consequences of range expansions. Annual Review of Ecology, Evolution, and Systematics, 40:481–501, 2009. doi: 10.1146/annurev.ecolsys.39.110707.173414.

[14] Adam Eyre-Walker and Peter D. Keightley. The distribution of fitness effects of new mutations. Nature Reviews Genetics, 8:610–618, 2007. doi: 10.1038/nrg2146.

[15] Julien Fayard, Etienne K. Klein, and François Lefèvre. Long distance dispersal and the fate of a gene from the colonization front. Journal of Evolutionary Biology, 22:2171–2182, 2009. doi: 10.1111/j.1420-9101.2009.01832.x.

[16] Joseph Felsenstein. The evolutionary advantage of recombination. Genetics, 78:737–756, 1974.

[17] Ronald A. Fisher. The wave of advance of advantageous genes. Annals of Eugenics, 7:355–369, 1937. doi: 10.1111/j.1469-1809.1937.tb02153.x.

[18] Jimmy Garnier, Thomas Giletti, François Hamel, and Lionel Roques. Inside dynamics of pulled and pushed fronts. Journal de Mathématiques Pures et Appliquées, 98:428–449, 2012. doi: 10.1016/j.matpur.2012.02.005.

[19] Kimberly J. Gilbert, Nathaniel P. Sharp, Amy L. Angert, Gina L. Conte, Jeremy A. Draghi, Frédéric Guillaume, Anna L. Hargreaves, Remi Matthey-Doret, and Michael C. Whitlock. Local adaptation interacts with expansion load during range expansion: Maladaptation reduces expansion load. The American Naturalist, 189:368–380, 2017. doi: 10.1086/690673.

[20] Kimberly J. Gilbert, Stephan Peischl, and Laurent Excoffier. Mutation load dynamics during environmentally-driven range shifts. PLOS Genetics, 14:1–18, 2018. doi: 10.1371/journal.pgen.1007450.

[21] Santiago C. González-Martínez, Kate Ridout, and John R. Pannell. Range expansion compromises adaptive evolution in an outcrossing plant. Current Biology, 27:2544–2551.e4, 2017. doi: 10.1016/j.cub.2017.07.007.

[22] Eva Graciá, Francisco Botella, José Daniel Anadón, Pim Edelaar, D. James Harris, and Andrés Giménez. Surfing in tortoises? Empirical signs of genetic structuring owing to range expansion. Biology Letters, 9, 2013. doi: 10.1098/rsbl.2012.1091.

[23] Matti Gralka, Fabian Stiewe, Fred Farrell, Wolfram Möbius, Bartlomiej Waclaw, and Oskar Hallatschek. Allele surfing promotes microbial adaptation from standing variation. Ecology Letters, 19:889–898, 2016. doi: 10.1111/ele.12625.

[24] Karl-Peter Hadeler and Franz Rothe. Travelling fronts in nonlinear diffusion equations. Journal of Mathematical Biology, 2:251–263, 1975. doi: 10.1007/BF00277154.

[25] John Haigh. The accumulation of deleterious genes in a population—Muller’s Ratchet. Theoretical Population Biology, 14:251–267, 1978. doi: 10.1016/0040-5809(78)90027-8.

[26] Oskar Hallatschek and David R. Nelson. Gene surfing in expanding populations. Theoretical Population Biology, 73:158–170, 2008. doi: 10.1016/j.tpb.2007.08.008.

[27] Oskar Hallatschek and David R. Nelson. Life at the front of an expanding population. Evolution, 64:193–206, 2010. doi: 10.1111/j.1558-5646.2009.00809.x.

[28] Oskar Hallatschek, Pascal Hersen, Sharad Ramanathan, and David R. Nelson. Genetic drift at expanding frontiers promotes gene segregation. Proceedings of the National Academy of Sciences, 104:19926–19930, 2007. doi: 10.1073/pnas.0710150104.

[29] Brenna M. Henn, L. Luca Cavalli-Sforza, and Marcus W. Feldman. The great human expansion. Proceedings of the National Academy of Sciences, 109:17758–17764, 2012. doi: 10.1073/pnas.1212380109.

[30] Kevin Higgins and Michael Lynch. Metapopulation extinction caused by mutation accumulation. Proceedings of the National Academy of Sciences of the United States of America, 98: 2928–2933, 2001. doi: 10.1073/pnas.031358898.

[31] Seraina Klopfstein, Mathias Currat, and Laurent Excoffier. The fate of mutations surfing on the wave of a range expansion. Molecular Biology and Evolution, 23:482–490, 2006. doi: 10.1093/molbev/msj057.

[32] Andrey Kolmogoroff, Ivan Petrovsky, and Nikolaj Piscounoff. Études de l’équation avec croissance de la quantité de matière et son application à un problème biologique. Moscow University Bulletin Of Mathematics, 1:1–25, 1937.

[33] Kirill S. Korolev, Melanie J. I. Müller, Nilay Karahan, Andrew W. Murray, Oskar Hallatschek, and David R. Nelson. Selective sweeps in growing microbial colonies. Physical Biology, 9: 026008, 2012. doi: 10.1088/1478-3975/9/2/026008.

[34] Andrew M. Kramer, Brian Dennis, Andrew M. Liebhold, and John M. Drake. The evidence for Allee effects. Population Ecology, 51:341, 2009. doi: 10.1007/s10144-009-0152-6.

[35] Laurence Loewe. Quantifying the genomic decay paradox due to Muller’s ratchet in human mitochondrial DNA. Genetical Research, 87:133–159, 2006. doi: 10.1017/S0016672306008123.

[36] Carl Müller and Roger Tribe. Stochastic P.D.E.’s arising from the long range contact and long range voter processes. Probability Theory and Related Fields, 102:519–545, 1995. doi: 10.1007/BF01198848.

[37] Hermann J. Muller. The relation of recombination to mutational advance. Mutation Research/Fundamental and Molecular Mechanisms of Mutagenesis, 1:2–9, 1964. doi: 10.1016/0027-5107(64)90047-8.

[38] Stephan Peischl and Laurent Excoffier. Expansion load: recessive mutations and the role of standing genetic variation. Molecular Ecology, 24:2084–2094, 2015. doi: 10.1111/mec.13154.

[39] Stephan Peischl, Isabelle Dupanloup, Mark Kirkpatrick, and Laurent Excoffier. On the accumulation of deleterious mutations during range expansions. Molecular Ecology, 22:5972–5982, 2013. doi: 10.1111/mec.12524.

[40] Stephan Peischl, Mark Kirkpatrick, and Laurent Excoffier. Expansion load and the evolutionary dynamics of a species range. The American naturalist, 185:E81–E93, 2015. doi: 10.1086/680220.

[41] Stephan Peischl, Isabelle Dupanloup, Lars Bosshard, and Laurent Excoffier. Genetic surfing in human populations: from genes to genomes. Current Opinion in Genetics & Development, 41:53–61, 2016. doi: 10.1016/j.gde.2016.08.003.

[42] Stephan Peischl, Isabelle Dupanloup, Adrien Foucal, Michèle Jomphe, Vanessa Bruat, JeanChristophe Grenier, Alexandre Gouy, Kimberly J. Gilbert, Elias Gbeha, Lars Bosshard, Elodie Hip-Ki, Mawussé Agbessi, Alan Hodgkinson, Hélène Vézina, Philip Awadalla, and Laurent Excoffier. Relaxed selection during a recent human expansion. Genetics, 208:763–777, 2018. doi: 10.1534/genetics.117.300551.

[43] Lionel Roques, Jimmy Garnier, François Hamel, and Etienne K. Klein. Allee effect promotes diversity in traveling waves of colonization. Proceedings of the National Academy of Sciences, 109:8828–8833, 2012. doi: 10.1073/pnas.1201695109.

[44] Yuval B. Simons, Michael C. Turchin, Jonathan K. Pritchard, and Guy Sella. The deleterious mutation load is insensitive to recent population history. Nature Genetics, 46:220–224, 2014. doi: 10.1038/ng.2896.

[45] Philip A. Stephens, William J. Sutherland, and Robert P. Freckleton. What is the Allee effect? Oikos, 87:185–190, 1999. doi: 10.2307/3547011.

[46] A. N. Stokes. On two types of moving front in quasilinear diffusion. Mathematical Biosciences, 31:307–315, 1976. doi: 10.1016/0025-5564(76)90087-0.

[47] Caz M. Taylor and Alan Hastings. Allee effects in biological invasions. Ecology Letters, 8: 895–908, 2005. doi: 10.1111/j.1461-0248.2005.00787.x.

[48] Justin M. J. Travis, Tamara Münkemüller, Olivia J. Burton, Alex Best, Calvin Dytham, and Karin Johst. Deleterious mutations can surf to high densities on the wave front of an expanding population. Molecular Biology and Evolution, 24:2334–2343, 2007. doi: 10.1093/molbev/msm167.

[49] Wim Van Saarloos. Front propagation into unstable states. Physics Report, 386:29–222, 2003. doi: 10.1016/j.physrep.2003.08.001.

[50] Yvonne Willi, Marco Fracassetti, Stefan Zoller, and Josh Van Buskirk. Accumulation of mutational load at the edges of a species range. Molecular Biology and Evolution, 35:781–791, 2018. doi: 10.1093/molbev/msy003.

